# Interactions among rooting traits for deep water and nitrogen uptake in upland and lowland ecotypes of switchgrass (*Panicum virgatum* L.)

**DOI:** 10.1101/2021.02.19.432036

**Authors:** Marcus Griffiths, Xueyan Wang, Kundan Dhakal, Haichao Guo, Anand Seethepalli, Yun Kang, Larry M. York

## Abstract

Plant phenotypic plasticity in response to nutrient and water availability is an important adaptation for abiotic stress tolerance. Roots intercept water and nutrients while foraging through soil searching for further resources. Substantial amounts of nitrate can leach into groundwater; yet, little is known about how deep rooting affects this process. Here, we phenotyped root system traits and deep ^15^N nitrate capture across 1.5 m profiles of solid-media using tall mesocosms in switchgrass (*Panicum virgatum* L.), a cellulosic bioenergy feedstock. Root and shoot biomass, photosynthesis and respiration, and nutrient uptake traits were quantified in response to a water and nitrate stress factorial experiment in the greenhouse for switchgrass upland (VS16) and lowland (AP13) ecotypes. The two switchgrass ecotypes shared common plastic abiotic responses to nitrogen (N) and water availability and yet showed genotypic differences for root and shoot traits. A significant interaction between nitrogen and water stress for axial and lateral root traits represents a complex and shared root development strategy for stress mitigation. Deep root growth and ^15^N capture were found to be closely linked to aboveground growth. Together, these results represent the wide genetic pool of switchgrass and that deep rooting promotes nitrate capture, plant productivity, and sustainability.

**Highlight:** Two main ecotypes of switchgrass have both shared and different root responses to varying water and nitrogen conditions, with deep rooting shown to be closely linked to aboveground growth.

## Introduction

The root system of a plant serves multiple important roles, from structural stability in the soil to resource foraging for water and nutrients. The spatial and temporal arrangement of roots in the soil (broadly referred to as root system architecture) can greatly affect the interception and subsequent uptake of soil resources. Root growth and development are highly responsive to both the environmental conditions and the plant’s resource requirements. Greater knowledge of this dynamic process in plants is important to characterize ecological adaptations and breed for beneficial adaptations enabling more resource-efficient plant varieties.

Water and nitrogen (N) are the two most frequently limiting resources in agriculture, affecting plant growth and yield. Both water and nitrate-N are highly mobile in the soil profile, which means plants have a limited opportunity to acquire these resources. Plant adaptive responses help mitigate such abiotic stresses through changes in growth and development in response to the plant’s nutritional requirements and the environment. Deep rooting is regarded as a beneficial trait for plant productivity and abiotic stress mitigation by expanding the soil volume explored, effectively increasing soil resources available to the plant. It increases the soil volume explored and consequently soil resources available to the plant (reviewed by Thorup-Kristensen et al., 2020). Increased root length at depth also enables plants to capture water and nitrate that otherwise would be lost through deep soil-water movement and leaching. Deep rooting traits in turn can reduce the environmental damage caused by the leaching of nutrients into groundwater (Foulkes et al., 2009; Kumar and Goh, 2002). However, roots are challenging to phenotype and evaluate quantitatively as they are the hidden half of the plant, and relatively little is known about root growth, nutrient capture, and root longevity of perennial crops. At present, root evaluation at depth is often conducted by soil coring methods in the field, with subsequent root washing and image analysis or qPCR for DNA abundance used for quantifying root length or mass (Kristensen and Thorup-Kristensen, 2004; Heuermann et al., 2019), stable isotope tracing (Ehleringer and Dawson, 1992; Chen et al., 2018), or under more controlled setups using large rhizotrons (Nagel et al., 2012; Ytting et al., 2014) or large mesocosms (Guo and York, 2019; Saengwilai et al., 2014; Zhan et al., 2015).

Switchgrass (*Panicum virgatum* L.) is a C4, warm-season perennial grass that is native to North America and has an extensive and deep root system with recorded rooting depths of 330 cm in field trials (Ma et al., 2000). As with many prairie grasses, switchgrass develops rhizomes which are underground, stem-derived organs that provide plants with the ability to grow clonally and regrow after disturbance in the soil (Weaver, 1954; Freschet et al., 2020). Switchgrass is found across a diverse geographical range from Canada to Central America and has a promising utility as a cellulosic bioenergy feedstock. Switchgrass has low input requirements, so is ideal for growth on marginal lands. In addition, switchgrass is reported to provide ecosystem services with an enhancement of soil organic matter, reduction in soil erosion, and associative N fixation (Lai et al., 2018; Gilley et al., 2000; Roley et al., 2019). Switchgrass can be divided into two main ecotypes, upland and lowland, which are estimated to have diverged 0.7–1.0 million years ago (Morris et al., 2011). The ecotype divergence in switchgrass is hypothesized to be through climatic-associated adaption with the upland ecotype found in more northern latitudes and across drier precipitation gradients than the lowland ecotype (Lovell et al., 2021). The upland ecotype has also been found to be generally smaller with a greater number of tillers and an earlier flowering time (Milano et al., 2016; Singer et al., 2019). As the ecotypes are diverse, each has its own beneficial breeding potential with different environmental adaptation and pathogen resistance (Milano et al., 2016). For switchgrass adoption as a bioenergy feedstock, the biomass yield will have to be maximized in a sustainable manner, which requires a greater understanding of the interactions among environment, ecotype, and soil dynamics (Lemus et al., 2014).

In-depth characterization of the physiological and morphological differences of the main switchgrass ecotypes is important to understand the functional adaptations to resource capture and to characterize the differences in abiotic stress tolerance. The aim of this study was to characterize and compare the root systems of the representative upland and lowland ecotypes of switchgrass and the root adaptive responses to water and nitrogen stress. To achieve this, we set up a tall mesocosm greenhouse study with water and nitrogen factorial stresses using clones of representative upland and lowland cultivars and evaluated the vertical distribution of the root system across 150 cm depth along with other physiological characteristics.

## Materials and methods

### Plant materials and experiment design

Clones derived from two contrasting genotypes of switchgrass, AP13 and VS16, were used in this study to represent the two ecotypes. AP13 is a clone derived from the lowland cultivar ‘Alamo’, which is the source of the switchgrass reference genome, and VS16 is a clone derived from the upland cultivar ‘Summer’. Mapping populations have been derived from crossing these two ecotypes (Milano et al., 2016), which highlights their importance for switchgrass research. Recently-emerged tillers from well-established plants were pulled apart by hand and one tiller consisting of a small shoot and root system was transplanted per mesocosm at the start of the experiment. The mesocosm experiment was conducted in a greenhouse from 30^th^ September 2019 to 22^nd^ January 2020 at the Noble Research Institute, LLC, Ardmore, OK, USA (34°11’ N, 97°5’ W; elevation 268 m). The greenhouse conditions were set to a 15/7 h day/night cycle at 24/21°C with an average photosynthetically active radiation (PAR) reading of 150 mol m^2^ s^1^ provided with supplemental lighting. Monthly averages for greenhouse conditions are provided in Table S1. The mesocosm experiment was arranged as a randomized complete block design, replicated five times with a 2×2×2 factorial arrangement of treatments. The factors were two levels of N supply (high- and low-N, HN and LN), two watering levels (well-watered and drought-stressed, WW and DS), and two ecotypes (upland and lowland). The treatment combinations are hereafter referred to as HN/WW, LN/WW, HN/DS, and LN/DS.

### Mesocosm preparation

The media mixture used in the mesocosm study mimics mineral soil and consisted of sand, vermiculite, and perlite which was mixed using a cement mixer. By volume basis, the mixture constituents used were 50% medium size (0.3–0.5 mm) premium sand (Quikrete, GA, USA), 40% premium grade vermiculite (Sun Gro Horticulture, Agawam, MA, USA), and 10% perlite (Ambient Minerals, AR, USA). The gravimetric water content of the media mixture at mesocosm filling and before watering was 2.5%, as determined by oven-drying 20 g of media at 105 °C for 48 hours (Equation 1) (Rowell, 1994).

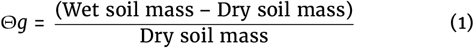

Media N levels of the starting media mixture before the nutrient application was determined to be 0.11 mM by ion chromatography. Twenty grams of media were first added to 50 mL of 0 N ½ Hoagland’s solution (detailed below) and then shaken for 30 min at 150 rpm. After shaking, the sample was left undisturbed for five minutes for the particles to settle, and then five mL of the supernatant was centrifuged at 10,000 rpm for five minutes in a 15 mL falcon tube. The ion concentrations of the collected supernatant samples were then determined using a Thermo Scientific ICS-5000+ ion chromatographic system using 500 L of the sample (Thermo Fisher Scientific, MA, USA).

The mesocosms used in this study consisted of polyvinyl chloride (PVC) pipe cut to size 15.24 cm [internal diameter] × 152.4 cm [height] with a flat-bottom PVC cap (IPS Corporation, Collierville, TN, USA), and lined with a seamless heavy-duty (6 Mil) poly tubing (Uline, WI, USA) (Fig. 1A). The mesocosms were evenly filled from the top of the column with 28 L of the air-dry media resulting in an approximate bulk density of 1.1 g cm^-3^. Three days before transplanting, the mesocosms were irrigated from the top with six L of nutrient solution. Half of the mesocosms received a zero N ½-strength Hoagland’s solution for LN treatment and the other received a high N ½-strength Hoagland’s solution (6 mM NO3-N) for HN treatment. The high N solution composed of (in µM) 500 KH_2_PO, 5700 KNO_3_, 300 NH_4_NO_3_, 2000 CaCl_2_, 1000 MgSO_4_, 46 H_3_BO_3_, 7 ZnSO_4_·_7_H_2_O, 9 MnCl_2_·_4_H_2_O, 0.32 CuSO_4_·_5_H_2_O, 0.114 (NH_4_)_6_Mo_7_O_24_·_4_H_2_O, and 150 Fe(III) EDTA(C_10_H_12_N_2_NaFeO_8_). For the zero N solution, KNO_3_ and NH_4_NO_3_ was replaced with 5700 KCl.

**Figure 1.**
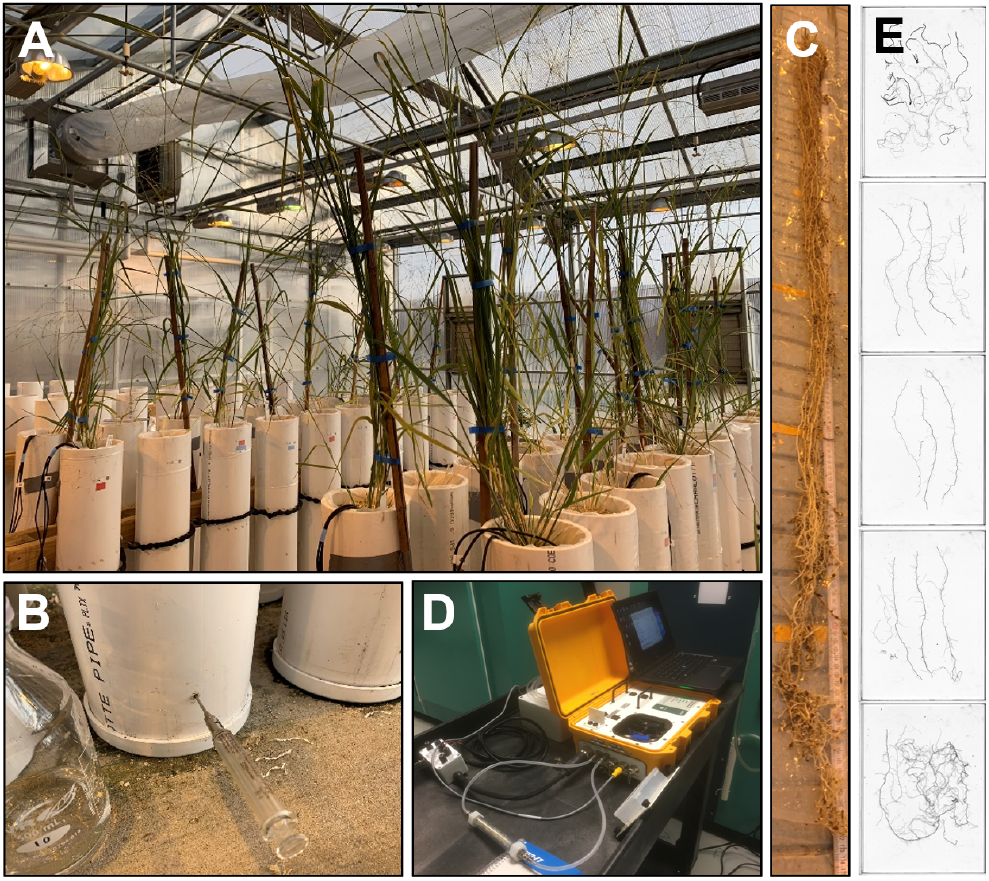
Switchgrass mesocosm experiment design. (A) Upland and lowland ecotypes were grown in tall mesocosms under a factorial nitrogen and water stress conditions HN/WW, LN/WW, HN/DS, LN/DS. (B) ^15^ N was injected into the deepest layer of the mesocosms 24 hours before the shoot material was harvested. (C) The medium was carefully washed away, and the root system was cut into 30 cm layers which were used for (D) instantaneous root respiration analysis using an LI-8100 with custom chambers, and (E) root feature determination by root scanning and image analysis using RhizoVision Explorer.

### Mesocosm greenhouse growth conditions

One tiller of a clone was transplanted per mesocosm; the mesocosms were watered with the respective nutrient solution three times a week with 200 mL added from the top at each watering. After four weeks of growth in the mesocosms, half of the mesocosms were subjected to drought-stress receiving no more watering for the rest of the experiment. The well-watered mesocosms continued to be watered three times a week with 200 mL double-deionized water instead of nutrient solution. At the end of the experiment, the drought-stressed mesocosms had a gravimetric water content ranging from 16% in the first 30 cm layer to 28% in the deepest 30 cm layer (Table 1). The gravimetric water content of the well-watered mesocosms ranged from 27% in the first 30 cm layer to 34% in the bottom layer (Table 1.). The water stress was applied over a depth gradient. Averaged across the whole mesocosm the gravimetric water content for the water-stressed and well-watered mesocosms were 23% and 29%, respectively.

**Table 1.** Gravimetric water content (%) of mesocosms determined at the end of the greenhouse study.

| Soil horizon<br>(cm depth) | Mesocosm treatment |  |
| --- | --- | --- |
|  | Well-watered | Drought-stressed |
| 0–30 | 27.28 | 16.47 |
| 30–60 | 26.52 | 23.26 |
| 60–90 | 27.36 | 19.80 |
| 90–120 | 29.84 | 26.04 |
| 120–150 | 33.86 | 27.86 |

### Mesocosm sample collection and harvest measures

Four months after the flowering onset, the plants were destructively harvested. Phenotypic traits measured are defined in Table 2. One day before destructive sampling, plant height was recorded using a ruler measured from the mesocosm media surface to the tip of the tallest leaf when held to its maximum height and all tillers were counted. Gas exchange and chlorophyll fluorescence parameters for the youngest fully expanded leaf of each plant were measured using an LI-6800 portable photosynthesis system with Multiphase Flash Fluorometer (LI-COR Biosciences, Lincoln, NE, USA) operating with a six cm^2^ aperture circular leaf adapter, a flow rate of 600 µmol mol^-1^, and a cuvette relative humidity of 60%. CO_2_ exchange was logged manually, with stability criteria for both 2O and 2 standard deviation limits set to 0.1 over a period of 15 s. The leaf maximum width was used to normalize measurements on leaf material smaller than the circular leaf adapter. Then, the mesocosms were injected with ^15^NO_3_ (98% atom) to assess deep N capture by switchgrass roots. Three evenly-spaced passage holes were drilled around the circumference of each mesocosm at 132 cm depth, and five mL of Ca(^15^ NO_3_)_2_ solution (0.46 mg ^15^NO_3_ mL^-1^) was injected into each mesocosm using these holes with a syringe (Fig. 1B).

**Table 2.**
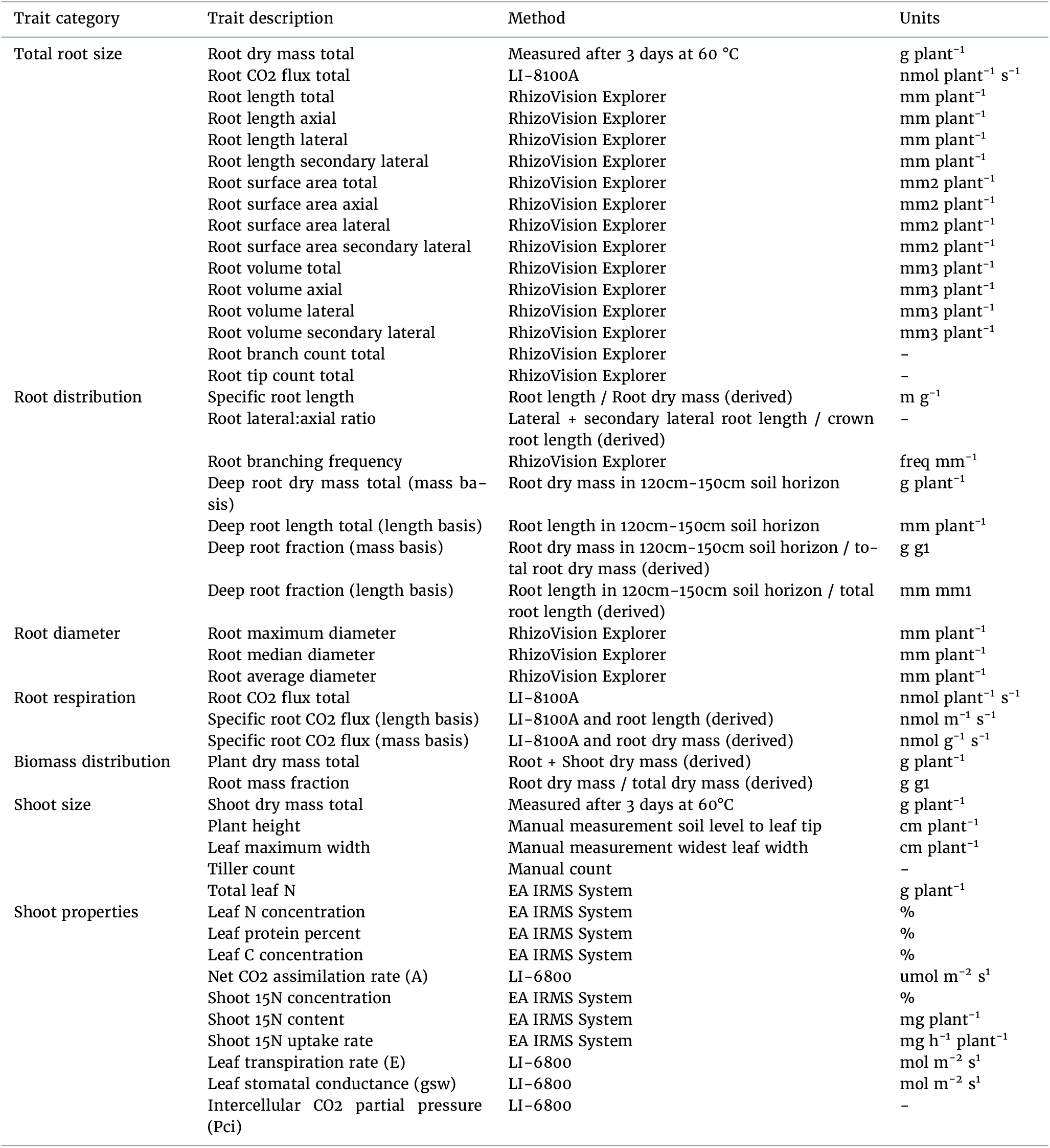
Traits measured and descriptors used in this study. Calculations for derived traits are found in the Supplementary R code. Traits measured in per plant basis refers to the entire plant within one biological unit.

The next day at 24 hours after ^15^N injection, the shoot of each plant was severed at the stem base and dried at 60 °C for 3 d for dry biomass determination. The shoot samples were thoroughly ground by placing the samples into glass vials with three opposing surgical blades and shaking at a frequency of 30 Hz for 10 minutes using a Qiagen TissueLyser II (Germantown, MD, USA). Shoot tissue percentages of total N and ^15^N were determined using a BioVisION from Elementar including an IsoPrime Vision isotope ratio mass spectrometer connected to an IsoPrime Isotope cube that operates in CNS mode (Elementar, Langenselbold, Germany).

For root harvesting, the polyethylene bag that lined each mesocosm was pulled out and the bag was sliced open longitudinally on a root washing station (Fig. 1C). One-hundred grams of media mixture samples excluding roots were bagged at 30 cm layers for measuring gravimetric water content and N content, as detailed above, and were placed in a cool box containing ice and frozen at -20 °C within eight hours. The rest of the mixture was then carefully washed away from the roots using a water hose with a low-pressure nozzle starting at the plant base. Immediately after root washing, roots were cut and divided into 30 cm layers, and root respiration for each plant and layer was measured (Fig. 1D and E). All roots from each layer were transferred into a custom 43 mL airtight chamber (as detailed in (Guo et al., 2020)) connected to an LI-8100 Automated Soil CO2 Flux System (LI-COR Biosciences, NE, USA). A representative subsample of roots was measured if there was too much root materials to fit into the chamber. The CO2 flux in the chamber was measured with an observation duration of 90 seconds using the LI-8100A v4.0.9 software (Fig. 1D). Total respiration rate was calculated automatically by the linear fit mode in SoilFluxPro v4.0.1 software (LI-COR Biosciences, NE, USA) with a curve-fit time of 20–90 seconds. After the root respiration measurement, the root material was bagged individually by plant, media layer, and by subsample if required (Fig. 1E). The root material was then placed in a cool box containing ice and frozen at -20 °C within eight hours.

The bagged root samples were later thawed and imaged using a flatbed scanner equipped with a transparency unit (Epson Expression 12000XL, Epson America Inc, Los Alamitos, CA, USA). Roots were spread out on a transparent acrylic tray (420 mm x 300 mm) with a five mm layer (400 mL) of water and imaged at a resolution of 600 dpi as JPG files with 95% (high) quality. Multiple root scans were done when too much root material was present to scan in a single image to minimize root overlapping based on subjective determination, with an average root length of 10 m per scan retroactively calculated, and the cumulative root length was computed in R. The axial, first-order lateral and second-order lateral root lengths, surface area, and volume for each plant was calculated from the flatbed images using RhizoVision Explorer v2.0.2 (Seethepalli and York, 2020) based on diameter thresholds (in mm) of > 0.9, 0.3–0.9, and < 0.3, respectively. The threshold level was set to 200, filter non-root objects set to 10 mm^2^, and root pruning threshold set to 20 pixels. Total root tip number, branching frequencies, and average root diameter were also calculated in the software. During statistical analysis, the ratio of lateral roots (first- and second-order) to the axial traits was computed as lateral-to-axial ratios. After scanning, the root material was placed in a paper envelope and dried at 60 °C for three days for determination of root dry weight. Root mass fraction was calculated by dividing the total root dry mass by the total plant dry mass, and deep root fraction as the root length or mass in the bottom 120–150 cm layer divided by the respective total root system length or mass.

## Statistical analysis

Statistical analyses were conducted using R version 4.0.3 (R Core Team, 2020); the statistical analysis R codes including the packages needed are available at https://doi.org/10.5281/zenodo.4281435 (Griffiths et al., 2021). Traits calculated are described in Table 2. Analysis of variance (ANOVA) of the plant data was conducted using the R package “lmerTest” (Kuznetsova et al., 2017) with block as the random effect. The Tukey’s HSD test used for the multiple comparison boxplots was conducted using the R package “agricolae” (De Mendiburu and Simon, 2015). The correlation matrix was generated using the R package “corrplot” (Wei and Simko, 2017) with plant trait data for both genotypes across all conditions. Linear discriminant analysis was conducted using the ‘lda’ function from the MASS package (Venables and Ripley, 2002) to predict genotype, water treatment, or nitrogen treatment classes in separate analyses. Before LDA, data were standardized for each trait so that the mean was zero and the within-group standard deviation was 1 in order to interpret loadings.

## Results

### Positive correlations among root size traits and deep rooting traits with shoot size traits in switchgrass

Across both switchgrass ecotypes and all conditions, a correlation matrix showed strong positive correlations among root size-related traits, deep rooting traits, shoot size-related traits, and ^15^N content of leaves (P < 0.05, Fig. 2). Root length, surface area, and volume traits were highly correlated (P < 0.05, Fig. S1). For the specific root traits, positive correlations were observed between specific root length, specific root respiration (length and mass basis), and ^15^N percent of leaves (P < 0.05, Fig. 2). For deep root fraction, the only correlated traits, aside from deep root mass, were for secondary lateral root traits and plant height (P < 0.05, Fig. 2). Gas exchange (Assimilation rate, transpiration rate, and stomatal conductance) and chlorophyll fluorescence parameters for the new fully expanded leaf were uncorrelated with all other measured plant traits (Fig. 2).

**Figure 2.**
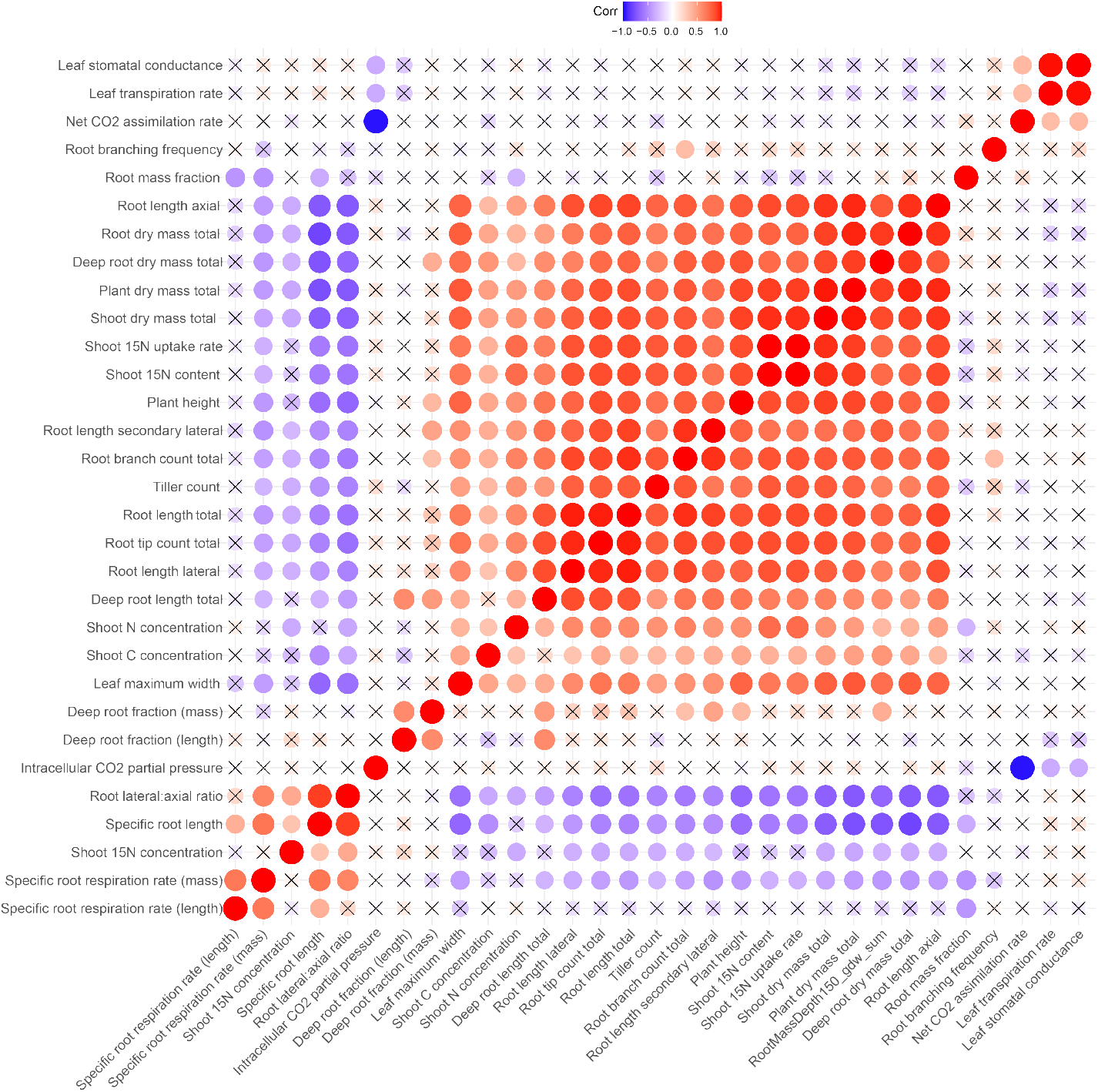
Correlation matrix for plant traits across both switchgrass ecotypes, upland (VS16) and lowland (AP13), and all conditions. Correlations are visualized using a color gradient. Red and blue color represent a strong positive and negative correlation respectively. No correlation is visualized with a cross symbol.

### Substantial phenotypic variation between the upland and lowland ecotypes for root and shoot traits representing the wide genetic pool of switchgrass

For the plant traits measured, substantial differences between the upland and lowland ecotypes were observed. Common to all conditions tested, total genotype-associated differences between the ecotypes were observed for total root mass, root class distribution traits including root mass fraction, specific root length, lateral:axial root ratio, and also specific root respiration rate (mass basis), plus leaf maximum width (P < 0.05, Fig. 3, Table S2). Across all conditions tested, the lowland ecotype had an average 80% larger root mass, 32% higher root mass fraction, 34% decrease in specific root length, 39% decrease in lateral:axial root ratio, and a 74% decrease in specific root respiration rate (mass basis) compared to the upland ecotype (Fig. 3, Table S2). In favorable conditions only, HN/WW, genotypic differences were also observed for tiller count with a 57% increase in the lowland (P < 0.05, Table S3 and S5). In all stress conditions tested (HN/DS, LN/WW, and LN/DS), genotypic differences were observed for root mass in the deepest mesocosm layer with a 50% larger root mass in the lowland relative to upland under high-N conditions and a 140% larger root mass in the upland compared to lowland under drought (P < 0.05, Table S4-S6). Under the most severe stress condition, LN/DS, genotypic differences were observed for axial root size traits with a 152% larger axial root system in the upland (length, surface area, volume) (P < 0.001, Table S4 and S6). Additional significant genotypic differences were observed in the LN/WW conditions for plant height, root branching frequency, root tip count, root length proportion in the deepest mesocosm layer compared to all layers, and total 15N content captured from the deepest layer with the upland being larger for all (P < 0.05, Table S6).

**Figure 3.**
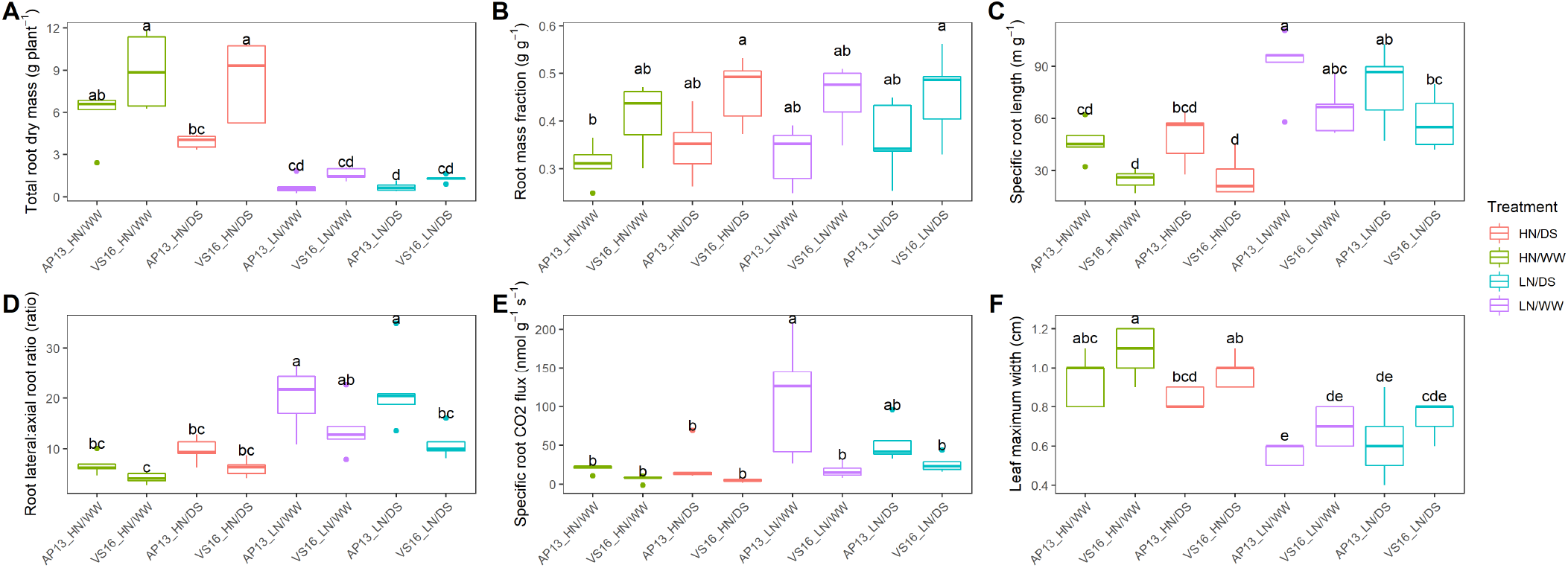
Total plant traits measured in each abiotic stress environment tested between the two switchgrass ecotypes, upland (VS16) and lowland (AP13). Boxes with the same letter were not significantly different at P < 0.05 according to Tukey’s HSD test.

A linear discriminant analysis using the plant traits was performed to determine the differentiating capacity of the plant traits between the two ecotypes. Across all conditions, rooting traits were the greatest discriminant trait between the ecotypes with axial and total root surface area being the greatest discriminators followed by root volume and length traits (Fig. 4A). In favorable conditions, HN/WW, the main discriminant traits were maximum tiller count, specific root length, leaf maximum width, specific root respiration rate (mass basis), and root mass fraction (Fig. 4B). Common to the well-watered conditions (HN/WW and LN/WW) was specific root respiration as a discriminant factor between the ecotypes (Fig. 4B and D). Common to the low-N conditions (LN/WW and LN/DS) was root maximum diameter as a discriminant factor (Fig. 4D and E). Specific to HN/DS, the main discriminant factors between the switchgrass ecotypes were for shoot N and 15N concentration, and plant height (Fig. 4C). Shoot 15N concentration was also a main discriminant for LN/WW. Specific to LN/WW, specific root respiration rate (both mass and length basis) and shoot carbon concentration were the main discriminant traits (Fig. 4D). Specific to LN/DS, root mass at depth and dry mass total were main the discriminant traits (Fig. 4E).

**Figure 4.**
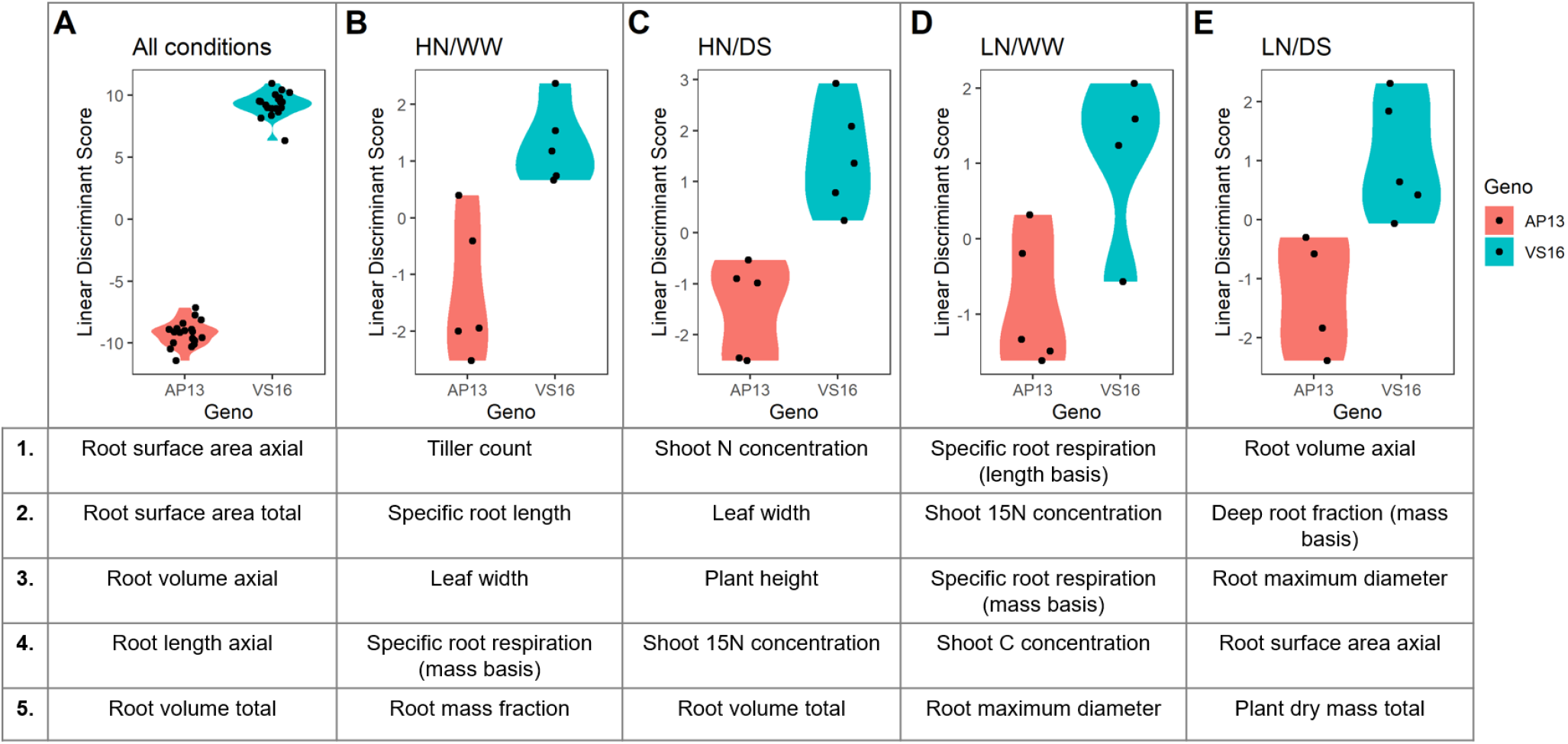
Linear discriminant analysis of total plant traits measured in each abiotic stress environment tested between the two switchgrass ecotypes, upland (VS16) and lowland (AP13). The five greatest discriminant traits by linear discriminant score both positive and negative are listed for each environment. (A) All conditions, (B) HN/WW, (C) HN/DS, (D) LN/WW, (E) LN/DS.

### Switchgrass ecotypes share common plastic abiotic responses to N and water availability

Genotypic differences in phenotypic traits were observed between the upland and lowland ecotypes, however, in terms of abiotic stress responses, common plastic responses were also observed between the ecotypes.

A significant water treatment response was common to both switchgrass ecotypes (HN conditions) with larger axial root traits (length, surface area, volume), lateral root traits (surface area, volume), root tip count, root maximum diameter, and shoot traits (tiller count, max leaf width) in well-watered conditions relative to drought-stressed conditions (P < 0.05, Table S5). A significant increase in lateral:axial root ratio and deep root fraction (mass basis) was observed in the water-stress conditions (HN conditions) (P < 0.01, Table S5). However, these water treatment effects were not observed in the LN conditions as the N stress appeared to have had a more severe and confounding effect on plant trait differences (Table S6).

In response to N treatment, a significant treatment effect was observed for all plant size traits with larger roots and shoots in high N conditions (WW and DS conditions, Table S3 and S4). In low N conditions, there was a significantly greater specific root length, lateral:axial root ratio, specific root respiration rate, and deep root fraction relative to the high N conditions. Traits with no significant difference by N treatment were for photosynthetic and transpiration measures (Table S3 and S4). No significant difference in root mass fraction was observed for either N treatments or water treatments with a shared reduction in both root and shoot mass by abiotic stress (Table S3 and S4). Under well-watered conditions, a significant genotype by N treatment interaction was observed for secondary lateral root size traits, branching frequency, and specific root respiration rate (mass basis) (P < 0.05, Table S3). Under drought conditions, a significant genotype by N treatment interaction was observed for root dry mass total, axial root size traits, deep root mass total, and deep root fraction (P < 0.05, Table S4).

A linear discriminant analysis using the plant traits was performed to determine the main discriminant trait between the water levels or N levels. The discriminant traits between N and W treatment levels were found to be rooting traits (Fig. 5A and B). For the N treatment, lateral and secondary lateral root traits were the top discriminant traits in addition to axial root surface area (Fig. 5A). For the W treatment, total root surface area and root surface area of each root class were the main discriminants plus axial root length (Fig. 5B).

**Figure 5.**
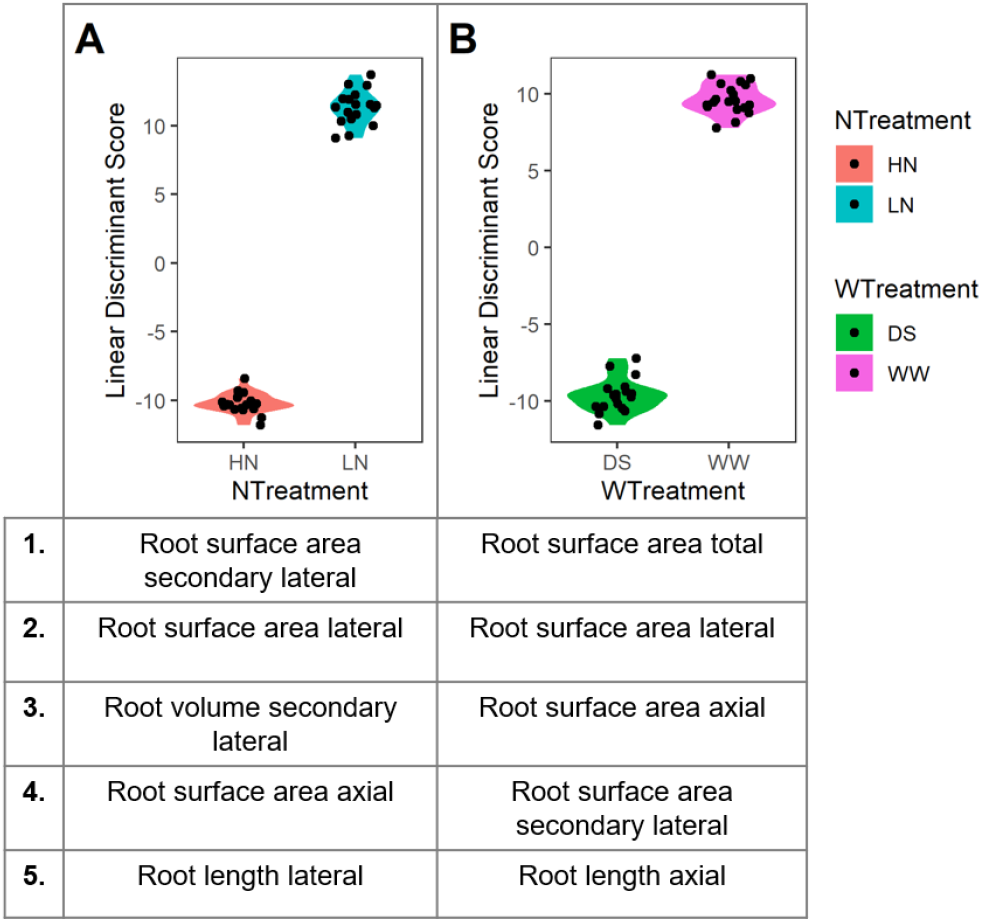
Linear discriminant analysis of total plant traits for both ecotypes to determine common discriminant traits by (A) N treatments, and (B) water treatments. The five greatest discriminant traits by linear discriminant score both positive and negative are listed from all environment data.

### Roots of both switchgrass ecotypes have the potential to grow deeper than 1.5 m with significant interaction between root depth related traits and abiotic stress

Across both ecotypes, almost all rooting traits tested were significantly affected by depth (Table S7-S10). Exceptions were for specific root respiration with no significant relationship with mesocosm depth and for lateral:axial root ratio in LN conditions.

Significant interaction between genotype and depth were observed for specific root respiration rate (weight basis) in favorable conditions, HN/WW. In the abiotic stress conditions, a significant genotype and depth interaction was observed for lateral root traits (length and surface area) in LN/WW, root axial traits (length and surface area) and root average diameter in LN/DS, and root maximum diameter in HN/DS.

The root distribution across the vertical profile varied greatly by water and nitrogen conditions (Fig. 6A). For both switchgrass ecotypes, total root length was greatest in the favorable condition, HN/WW, and least in the combined stress condition, LN/DS. In favorable conditions, there was no significant genotypic difference in root length by layer. However, in the low-N conditions, the upland ecotype had a greater root length in the deepest layer compared to the lowland ecotype. The greatest difference between the ecotypes was observed in the LN/WW condition with the upland ecotype having a 193% increased root length in the deepest layer, with the LN/DS condition reducing further the root length at depth for both ecotypes. The differences observed in the root length by depth also conferred with the shoot 15N content results with a 500% average greater 15N uptake in the HN condition plants than in the LN conditions, reflecting uptake 24 hours after 15N was injected to the bottom layer (Fig. 6B and C). An increase in lateral and secondary lateral roots in the upland ecotype contributed to this root length increase in the deepest mesocosm layer (Fig. 6A). A positive significant relationship was observed between the root length in the deepest layer and 15N content in shoot material with greater root length conforming to greater 15N uptake (P < 0.001, Fig. 6D).

**Figure 6.**
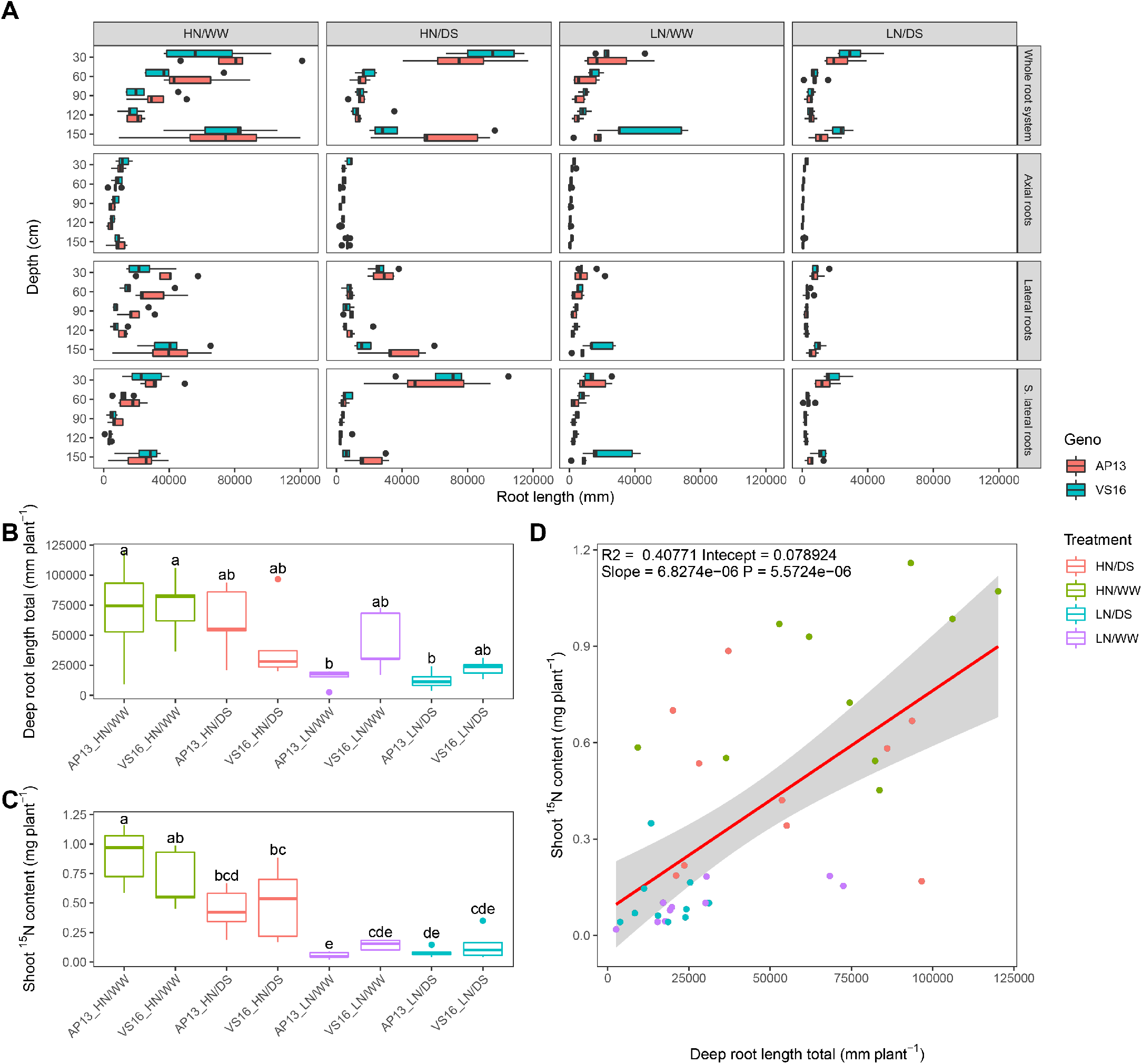
Root distribution of upland (VS16) and lowland (AP13) switchgrass ecotypes across 1.5 m mesocosms under abiotic stress environment. The roots distributions by root class were separated into 30 cm mesocosm layers. (B) Root length in the deepest layer and (C) 15N content in the shoot for the switchgrass ecotypes by treatment condition. Boxes with different letters were significantly different at P < 0.05 according to Tukey’s HSD test. (D) Linear regression analysis using all data between root length in the deepest layer and 15N content in the shoot.

### Significant interaction between N and W stress combination treatments for axial and lateral root traits

Using the whole dataset containing both ecotypes and all conditions, interactions between N and water treatment were explored. Across all conditions, significant interactions between N and water treatment were observed for axial root traits (length, surface area, volume), total root traits (volume, surface area), deep root fraction (mass basis) compared to all depths, shoot ^15^N content, ^15^N uptake rate, and shoot mass (P < 0.05, Table S11). In the lowland ecotype only, an interaction between N and water treatment was also observed for tiller count, total plant dry mass, and shoot N% (P < 0.05, Table S12). In the upland ecotype only, an interaction between N and water treatment was observed for secondary lateral root traits (length and surface area) (P < 0.05, Table S13).

## Discussion

Here, we show that members of two main ecotypes of switchgrass, upland (VS16) and lowland (AP13), share common root plastic response strategies to abiotic stress despite having large intrinsic root morphological differences. Appropriate growth responses to abiotic stress can be important stress mitigation strategies with efficient soil exploration for required resources. Despite previous studies finding switchgrass productivity of cultivar ‘Cave-in-Rock’ to be not receptive to fertilizer treatments (Duran et al., 2016), here a large N treatment effect was found in both switchgrass ecotypes. Potentially, this cultivar could have a different N response, the field experiment could have been growth-limited by other factors, or else the field soil had more residual available N than in the LN mesocosms. In response to N application in the current experiment, all plant size traits were significantly affected with an overall reduction in root and shoot size traits under stress conditions. Similarly, water-stress conditions in this study had significant plant size reducing effects for both root and shoot traits.

The switchgrass root system makes up a large proportion of the plant biomass with 34% of total biomass as roots for the lowland ecotype and 44% of total biomass in the upland ecotype in these single year plants, averaged across all treatments. Switchgrass can sequester a large amount of carbon and has been shown to increase soil carbon levels over time (Ma et al., 2001). Interestingly, the differences observed in root mass fraction in this study was by ecotype only with a stable fraction across water and N conditions. A significant interaction between drought and N stress conditions was observed in switchgrass for axial and lateral root traits representing a complex and shared root development strategy for stress mitigation. Both ecotypes had a smaller axial and lateral root length in the stressed conditions compared to the favorable conditions, probably driven by a reduction in growth and photosynthate availability. A similar relationship was found across 12 temperate herbaceous species with changes in belowground biomass allocation in response to nutrient supply but no change in root mass fraction (Freschet et al., 2015). For switchgrass, the main discriminants between favorable and stress conditions, water and N, across both ecotypes were for total root surface area size traits. Roots are a large carbon investment and maintenance cost to the plant and therefore a reduced root axial investment under stressed conditions is an efficient plastic root response. These innate responses to abiotic stress reflect the hardiness of the species and the ideal nature of switchgrass as a low-input crop (Vogel, 1996).

Specifically, in response to N conditions, lateral root traits were also found to be the main discriminants. Lateral roots are regarded as the primary site for the uptake of soil resources and often the greatest contributor to total root length and surface area in contact with soil (Hund et al., 2009; Yu et al., 2019). Total root length was reduced under N stress indicating that the plant was unable to sustain the total carbon cost of a large root system However, the inability to maintain the larger total size was partially compensated by more efficient carbon use by increasing the allocation to cheaper lateral roots, as shown by the greater lateral:axial root ratio. An increase in resource distribution to lateral roots, therefore, increased the root exploration in the soil with a reduced resource allocation to roots which can be seen as an efficient abiotic stress adaptive response, as also shown in maize (Guo and York, 2019). This highlights the importance of lateral roots for abiotic stress mitigation in switchgrass and that selection for improved, resource-efficient switchgrass varieties could use lateral:axial root ratio as a selection criterion for further investigation. This trait is convenient as it can be measured in a subsample of the root system, rather than requiring full excavation or measurements of entire root systems.

Switchgrass can be found across a wide range of climatic conditions and the upland and lowland ecotype represent the main divergent groups. Members of each ecotype, AP13 and VS16, were chosen for this study as AP13 is the source of the lowland reference genome and mapping populations have been derived from the two ecotypes (Milano et al., 2016). Variation among these ecotypes and others could be harnessed for improving abiotic stress tolerance and yield. Between these two switchgrass ecotypes, large morphological differences were observed in root traits with potential implications for abiotic stress tolerance. In a previous study, the upland ecotype was found to be more drought tolerant and had higher nitrogen demand than the lowland ecotype in 1-gallon pot trials, but the root traits were not quantified (Milano et al., 2016). In this single year study, a greater root mass was found in the upland ecotype in all conditions, which may be a contributor to its greater drought tolerance potential, although a shoot biomass difference was not observed. Across all conditions, rooting traits were the greatest discriminant between the ecotypes with lateral and secondary lateral roots being the main discriminants by N condition. The two ecotypes did not differ by shoot mass per condition in this study, but there was a significant tiller count difference with the lowland having a greater number of tillers in favorable conditions (HN/WW). Tiller counts in switchgrass have been shown to vary greatly year by year in field trials which is a likely response to the environment including rainfall patterns and competition with neighboring plants (Cassida et al., 2005; Price and Casler, 2014). The upland ecotype maintained the same number of tillers between the nitrogen and water conditions indicating stability across abiotic stress. The lowland ecotype tillered more in favorable conditions which may translate to an increase in overall shoot biomass and resilience across multiple years. Therefore, the upland and lowland ecotypes have varying strategies and adaptations that may translate to stress resistance and productivity in varying environments. Upland alleles have been previously associated with shoot size and vigor which may explain the greater root mass differences observed between the ecotypes in this study (Lowry et al., 2019).

An important plant trait for water and nutrient capture is deep rooting, however, it is technically challenging to excavate a representative root system from the field and quantify root length by soil depths. In this study, 1.5 m mesocosms were used to phenotype root distribution in switchgrass in 30 cm layers along the vertical profile with minimal root loss compared to field studies. Switchgrass is a particularly deep-rooted species, and in this study, a positive correlation was found between root length at depth and deep 15N capture by roots. Both ecotypes had roots in the deepest layer and had the potential to grow deeper than 1.5 m, given the substantial root length density in the bottom layers. Deep rooting is an important trait for crop performance because water and nitrate are often found in deep soil layers. Variance observed in switchgrass root size traits and 15N capture were found to explain differences in shoot mass highlighting the link between root and shoot. Between the upland and lowland ecotypes, differences in abiotic stress mitigation strategies were observed. Across all stress conditions tested, a genotype-associated difference was observed between the upland and lowland ecotypes for root mass in the deepest layer of the mesocosm. In the low-N conditions, a greater root length and mass were observed in the upland ecotype which conformed to a greater 15N shoot content which was applied to the deepest layer. Therefore, the upland ecotype was more receptive to the vertical N stress gradient with greater root development at depth and greater N uptake, which is an advantageous trait for low input cropping systems. Given the difficulties in excavating root systems and soil coring in the field, injection of ^15^N in deep layers and measuring uptake in the shoot is a viable method to screen for deep rooting activity. Interestingly, in response to drought conditions, the lowland ecotype had a smaller root mass compared to the upland ecotype in the deepest layer, however, the upland had a greater proportion of roots in this bottom layer. This indicates a root length distribution change to the vertical gradient water stress in the lowland ecotype and could be an advantageous drought tolerance trait.

Our findings highlight the importance of the root system with switchgrass ecotypes sharing common strategies for abiotic stress mitigation and deep N capture. We also show that the ecotypes have differing strategies to abiotic stress tolerance with biomass distribution changes and deep rooting in response to factorial water and N stress. Admixture between the divergent genomes is expected to enhance climate adaptation and yield improvement (Lovell et al., 2021). For switchgrass to be a productive bioenergy crop a balance between productivity and resource sustainability will have to be reached by enhancing plant abiotic stress tolerance and soil resource use efficiency.

## Availability of supporting data and materials

The dataset supporting the results of this article is available online as a Zenodo repository https://doi.org/10.5281/zenodo.4281435 (Griffiths et al., 2021).

## Author’s Contributions

L.M.Y. and M.G. conceived the research. M.G., H.G., K.D., and A.S. contributed to the experimentation. M.G., A.S., and L.M.Y. analyzed the data. All authors contributed to the writing of the manuscript.

## Acknowledgements

The authors would like to thank Michael Cloyde, Yaxin Ge, Amy Lee. and Jenifer Bonner for mesocosm sampling assistance; David Huhman and Bonnie Watson for operating the elemental analysis; and Jennifer Black and Malay Saha for providing the clonal material used in this study.

