## Supplemental Information for "Interactions among rooting traits for deep water and nitrogen uptake in upland and lowland ecotypes of switchgrass (*Panicum virgatum* L.)"

Table S1. Monthly average greenhouse conditions for study

| Month | Average monthly temperature (°C) | | Average monthly PAR (μmol m^−2^ s^−1^) |
| --- | --- | --- | --- |
|  | Day (15h) | Night (9h) |  |
| October | 24.87 | 19.92 | 151.06 |
| November | 24.22 | 18.60 | 141.40 |
| December | 23.83 | 18.22 | 150.02 |
| January | 23.56 | 18.27 | 144.60 |

Table S2. Genotype (all conditions)

|  | Traits | Source of variation |
| --- | --- | --- |
|  |  | Geno |
| Total root size | Root dry mass total (g plant^-1^) | 4.25 * |
|  | Root length total (mm plant^-1^) | 0.05 ns |
|  | Root length axial (mm plant^-1^) | 1.50 ns |
|  | Root length lateral (mm plant^-1^) | 0.25 ns |
|  | Root length secondary lateral (mm plant^-1^) | 0.65 ns |
|  | Root surface area total (mm^2^ plant^-1^) | 0.21 ns |
|  | Root surface area axial (mm^2^ plant^-1^) | 2.09 ns |
|  | Root surface area lateral (mm^2^ plant^-1^) | 0.42 ns |
|  | Root surface area secondary lateral (mm^2^ plant^-1^) | 0.66 ns |
|  | Root volume total (mm^3^ plant^-1^) | 1.06 ns |
|  | Root volume axial (mm^3^ plant^-1^) | 2.54 ns |
|  | Root volume lateral (mm^3^ plant^-1^) | 0.59 ns |
|  | Root volume secondary lateral (mm^3^ plant^-1^) | 0.66 ns |
|  | Root branch count total | 0.00 ns |
|  | Root tip count total | 0.03 ns |
| Root distribution | Specific root length (m g^-1^) | 8.83 ** |
|  | Root lateral:axial root ratio (ratio) | 6.96 * |
|  | Root branching frequency (branch mm^-1^) | 0.73 ns |
|  | Deep root mass total (g plant^-1^) | 2.82 ns |
|  | Deep root length total (mm plant^-1^) | 0.29 ns |
|  | Deep root mass fraction (g g^1^) | 0.80 ns |
|  | Deep root length fraction (mm mm^1^) | 2.60 ns |
| Root diameter | Root diameter mean (mm plant^-1^) | 4.29 * |
|  | Root diameter maximum (mm plant^-1^) | 0.10 ns |
|  | Root diameter median (mm plant^-1^) | 3.89 ns |
| Root respiration | Root CO_2_ flux total (nmol plant^-1^ s^-1^) | 8.24 ** |
|  | Specific root CO_2_ flux (nmol g^-1^ s^-1^) | 8.68 ** |
|  | Specific root CO_2_ flux (nmol m^-1^ s^-1^) | 4.47 * |
| Biomass distribution | Root mass fraction (g g^1^) | 31.50 *** |
|  | Total plant mass (g plant^-1^) | 1.28 ns |
| Shoot size | Shoot dry mass total (g plant^-1^) | 0.25 ns |
|  | Plant height (cm plant^-1^) | 0.81 ns |
|  | Tiller count | 1.80 ns |
|  | Leaf maximum width (cm) | 4.69 * |
| Shoot properties | Leaf carbon content (%) | 1.78 ns |
|  | Leaf N concentration (%) | 2.07 ns |
|  | Leaf 15N concentration (%) | 1.09 ns |
|  | Leaf total 15N content (mg plant^-1^) | 0.00 ns |
|  | Leaf 15N uptake rate (mg plant^-1^ h^-1^) | 0.00 ns |
|  | CO_2_ assimilation rate (µmol m^-2^ s^-1^) | 0.98 ns |
|  | Transpiration rate (mol m^-2^ s^-1^) | 0.02 ns |
|  | Stomatal conductance (mol m^-2^ s^-1^) | 0.01 ns |
|  | Intracellular CO_2_ (Pci) | 0.58 ns |

*** P < 0.001; ** P < 0.01; * P < 0.05; ns not significant

Table S3. Genotype x N condition (well-watered conditions)

|  | Traits | Source of variation | | |
| --- | --- | --- | --- | --- |
|  |  | Geno | N Treatment | Geno:N Treatment |
| Total root size | Root dry mass total (g plant^-1^) | 7.31 * | 69.17 *** | 2.40 ns |
|  | Root length total (mm plant^-1^) | 0.01 ns | 35.47 *** | 2.55 ns |
|  | Root length axial (mm plant^-1^) | 1.66 ns | 62.92 *** | 0.16 ns |
|  | Root length lateral (mm plant^-1^) | 0.39 ns | 31.93 *** | 2.63 ns |
|  | Root length secondary lateral (mm plant^-1^) | 0.59 ns | 17.62 *** | 4.89 * |
|  | Root surface area total (mm^2^ plant^-1^) | 0.15 ns | 53.14 *** | 0.69 ns |
|  | Root surface area axial (mm^2^ plant^-1^) | 3.06 ns | 76.61 *** | 0.78 ns |
|  | Root surface area lateral (mm^2^ plant^-1^) | 0.58 ns | 31.93 *** | 2.78 ns |
|  | Root surface area secondary lateral (mm^2^ plant^-1^) | 0.59 ns | 18.85 *** | 4.55 * |
|  | Root volume total (mm^3^ plant^-1^) | 1.50 ns | 78.42 *** | 0.08 ns |
|  | Root volume axial (mm^3^ plant^-1^) | 4.32 ns | 85.96 *** | 1.68 ns |
|  | Root volume lateral (mm^3^ plant^-1^) | 0.76 ns | 31.54 *** | 2.90 ns |
|  | Root volume secondary lateral (mm^3^ plant^-1^) | 0.57 ns | 19.91 *** | 4.27 ns |
|  | Root branch count total | 0.45 ns | 25.89 *** | 4.02 ns |
|  | Root tip count total | 0.01 ns | 72.03 *** | 4.19 ns |
| Root distribution | Specific root length (m g^-1^) | 20.87 *** | 65.28 *** | 0.14 ns |
|  | Root lateral:axial root ratio (ratio) | 5.85 * | 40.26 *** | 0.96 ns |
|  | Root branching frequency (branch mm^-1^) | 0.05 ns | 0.06 ns | 6.05 * |
|  | Deep root mass total (g plant^-1^) | 3.39 ns | 44.87 *** | 0.02 ns |
|  | Deep root length total (mm plant^-1^) | 1.73 ns | 11.67 ** | 0.97 ns |
|  | Deep root mass fraction (g g^1^) | 1.16 ns | 0.15 ns | 1.80 ns |
|  | Deep root length fraction (mm mm^1^) | 4.87 * | 0.21 ns | 0.13 ns |
| Root diameter | Root diameter mean (mm plant^-1^) | 9.84 ** | 113.07 *** | 2.58 ns |
|  | Root diameter maximum (mm plant^-1^) | 4.56 ns | 52.39 *** | 5.38 * |
|  | Root diameter median (mm plant^-1^) | 0.13 ns | 125.29 *** | 0.22 ns |
| Root respiration | Root CO_2_ flux total (nmol plant^-1^ s^-1^) | 19.10 ** | 21.07 *** | 3.33 ns |
|  | Specific root CO_2_ flux (nmol g^-1^ s^-1^) | 8.43 * | 7.59 * | 4.73 * |
|  | Specific root CO_2_ flux (nmol m^-1^ s^-1^) | 4.05 ns | 1.10 ns | 2.04 ns |
| Biomass distribution | Root mass fraction (g g^1^) | 16.01 ** | 1.18 ns | 0.20 ns |
|  | Total plant mass (g plant^-1^) | 2.22 ns | 105.19 *** | 0.45 ns |
| Shoot size | Shoot dry mass total (g plant^-1^) | 0.20 ns | 101.06 *** | 0.00 ns |
|  | Plant height (cm plant^-1^) | 1.31 ns | 155.83 *** | 7.59 * |
|  | Tiller count | 6.63 * | 34.61 *** | 3.38 ns |
|  | Leaf maximum width (cm) | 12.38 ** | 91.20 *** | 0.00 ns |
| Shoot properties | Leaf carbon content (%) | 1.11 ns | 46.12 *** | 1.79 ns |
|  | Leaf N concentration (%) | 2.85 ns | 24.57 *** | 5.40 * |
|  | Leaf 15N concentration (%) | 6.02 * | 7.28 * | 5.95 * |
|  | Leaf total 15N content (mg plant^-1^) | 0.59 ns | 81.88 *** | 3.79 ns |
|  | Leaf 15N uptake rate (mg plant^-1^ h^-1^) | 0.59 ns | 81.88 *** | 3.79 ns |
|  | CO_2_ assimilation rate (µmol m^-2^ s^-1^) | 0.39 ns | 0.30 ns | 0.49 ns |
|  | Transpiration rate (mol m^-2^ s^-1^) | 0.84 ns | 0.64 ns | 2.85 ns |
|  | Stomatal conductance (mol m^-2^ s^-1^) | 1.17 ns | 0.46 ns | 2.88 ns |
|  | Intracellular CO_2_ (Pci) | 0.30 ns | 0.31 ns | 0.15 ns |

*** P < 0.001; ** P < 0.01; * P < 0.05; ns not significant

Table S4. Genotype x N condition (droughted conditions)

|  | Traits | Source of variation | | |
| --- | --- | --- | --- | --- |
|  |  | Geno | N Treatment | Geno:N Treatment |
| Total root size | Root dry mass total (g plant^-1^) | 15.24 ** | 65.39 *** | 8.85 * |
|  | Root length total (mm plant^-1^) | 0.42 ns | 60.89 *** | 0.24 ns |
|  | Root length axial (mm plant^-1^) | 24.57 *** | 180.80 *** | 5.43 * |
|  | Root length lateral (mm plant^-1^) | 0.42 ns | 40.14 *** | 1.52 ns |
|  | Root length secondary lateral (mm plant^-1^) | 1.01 ns | 32.96 *** | 0.00 ns |
|  | Root surface area total (mm^2^ plant^-1^) | 2.53 ns | 125.47 *** | 0.00 ns |
|  | Root surface area axial (mm^2^ plant^-1^) | 31.27 *** | 177.18 *** | 11.09 ** |
|  | Root surface area lateral (mm^2^ plant^-1^) | 0.91 ns | 44.15 *** | 2.24 ns |
|  | Root surface area secondary lateral (mm^2^ plant^-1^) | 1.09 ns | 33.86 *** | 0.02 ns |
|  | Root volume total (mm^3^ plant^-1^) | 20.73 *** | 236.90 *** | 6.46 * |
|  | Root volume axial (mm^3^ plant^-1^) | 29.51 *** | 136.14 *** | 14.01 ** |
|  | Root volume lateral (mm^3^ plant^-1^) | 1.56 ns | 48.61 *** | 3.21 ns |
|  | Root volume secondary lateral (mm^3^ plant^-1^) | 1.15 ns | 34.27 *** | 0.04 ns |
|  | Root branch count total | 0.74 ns | 42.15 *** | 0.01 ns |
|  | Root tip count total | 0.19 ns | 52.23 *** | 2.15 ns |
| Root distribution | Specific root length (m g^-1^) | 8.32 * | 16.88 *** | 0.03 ns |
|  | Root lateral:axial root ratio (ratio) | 12.46 ** | 17.24 *** | 3.12 ns |
|  | Root branching frequency (branch mm^-1^) | 1.83 ns | 0.00 ns | 0.03 ns |
|  | Deep root mass total (g plant^-1^) | 11.10 ** | 87.74 *** | 5.39 * |
|  | Deep root length total (mm plant^-1^) | 0.31 ns | 11.99 ** | 2.44 ns |
|  | Deep root mass fraction (g g^1^) | 0.09 ns | 8.96 ** | 2.46 ns |
|  | Deep root length fraction (mm mm^1^) | 0.08 ns | 0.09 ns | 5.39 * |
| Root diameter | Root diameter mean (mm plant^-1^) | 17.25 ** | 82.15 *** | 0.41 ns |
|  | Root diameter maximum (mm plant^-1^) | 20.18 *** | 54.57 *** | 0.90 ns |
|  | Root diameter median (mm plant^-1^) | 0.25 ns | 26.80 *** | 0.00 ns |
| Root respiration | Root CO_2_ flux total (nmol plant^-1^ s^-1^) | 4.16 ns | 2.63 ns | 2.98 ns |
|  | Specific root CO_2_ flux (nmol g^-1^ s^-1^) | 6.86 * | 8.09 * | 0.17 ns |
|  | Specific root CO_2_ flux (nmol m^-1^ s^-1^) | 0.66 ns | 0.05 ns | 1.20 ns |
| Biomass distribution | Root mass fraction (g g^1^) | 13.77 ** | 0.01 ns | 0.15 ns |
|  | Total plant mass (g plant^-1^) | 6.41 * | 71.82 *** | 3.57 ns |
| Shoot size | Shoot dry mass total (g plant^-1^) | 1.64 ns | 57.43 *** | 0.84 ns |
|  | Plant height (cm plant^-1^) | 2.10 ns | 27.21 *** | 0.01 ns |
|  | Tiller count | 0.04 ns | 57.85 *** | 0.04 ns |
|  | Leaf maximum width (cm) | 8.59 * | 26.90 *** | 0.05 ns |
| Shoot properties | Leaf carbon content (%) | 4.92 * | 30.35 *** | 2.10 ns |
|  | Leaf N concentration (%) | 1.05 ns | 3.67 ns | 0.07 ns |
|  | Leaf 15N concentration (%) | 0.07 ns | 4.92 * | 0.07 ns |
|  | Leaf total 15N content (mg plant^-1^) | 0.52 ns | 17.37 *** | 0.00 ns |
|  | Leaf 15N uptake rate (mg plant^-1^ h^-1^) | 0.52 ns | 17.37 *** | 0.00 ns |
|  | CO_2_ assimilation rate (µmol m^-2^ s^-1^) | 0.54 ns | 0.00 ns | 1.46 ns |
|  | Transpiration rate (mol m^-2^ s^-1^) | 1.16 ns | 0.26 ns | 0.00 ns |
|  | Stomatal conductance (mol m^-2^ s^-1^) | 1.45 ns | 0.34 ns | 0.00 ns |
|  | Intracellular CO_2_ (Pci) | 0.25 ns | 0.00 ns | 0.75 ns |

*** P < 0.001; ** P < 0.01; * P < 0.05; ns not significant

Table S5. Genotype x W condition (high N conditions)

|  | Traits | | Source of variation | | |
| --- | --- | --- | --- | --- | --- |
|  |  | | Geno | W Treatment | Geno:W Treatment |
| Total root size | | Root dry mass total (g plant^-1^) | 14.99 ** | 1.72 ns | 0.37 ns |
|  | | Root length total (mm plant^-1^) | 0.41 ns | 2.26 ns | 0.54 ns |
|  | | Root length axial (mm plant^-1^) | 3.83 ns | 13.63 ** | 0.11 ns |
|  | | Root length lateral (mm plant^-1^) | 2.24 ns | 5.28 * | 0.28 ns |
|  | | Root length secondary lateral (mm plant^-1^) | 0.01 ns | 1.20 ns | 0.90 ns |
|  | | Root surface area total (mm^2^ plant^-1^) | 0.01 ns | 9.14 ** | 0.29 ns |
|  | | Root surface area axial (mm^2^ plant^-1^) | 7.58 * | 16.57 *** | 0.06 ns |
|  | | Root surface area lateral (mm^2^ plant^-1^) | 2.88 ns | 5.61 * | 0.26 ns |
|  | | Root surface area secondary lateral (mm^2^ plant^-1^) | 0.03 ns | 0.16 ns | 0.86 ns |
|  | | Root volume total (mm^3^ plant^-1^) | 3.30 ns | 17.29 *** | 0.13 ns |
|  | | Root volume axial (mm^3^ plant^-1^) | 11.18 ** | 18.53 *** | 0.04 ns |
|  | | Root volume lateral (mm^3^ plant^-1^) | 3.49 ns | 5.90 * | 0.25 ns |
|  | | Root volume secondary lateral (mm^3^ plant^-1^) | 0.04 ns | 0.01 ns | 0.81 ns |
|  | | Root branch count total | 0.60 ns | 0.01 ns | 1.62 ns |
|  | | Root tip count total | 2.30 ns | 5.45 * | 0.04 ns |
| Root distribution | | Specific root length (m g^-1^) | 22.31 *** | 0.18 ns | 0.00 ns |
|  | | Root lateral:axial root ratio (ratio) | 17.04 ** | 11.25 ** | 0.40 ns |
|  | | Root branching frequency (branch mm^-1^) | 0.14 ns | 0.29 ns | 3.19 ns |
|  | | Deep root mass total (g plant^-1^) | 7.28 * | 1.03 ns | 1.37 ns |
|  | | Deep root length total (mm plant^-1^) | 0.32 ns | 1.96 ns | 0.72 ns |
|  | | Deep root mass fraction (g g^1^) | 0.66 ns | 8.87 ** | 0.27 ns |
|  | | Deep root length fraction (mm mm^1^) | 0.00 ns | 0.76 ns | 4.23 ns |
| Root diameter | | Root diameter mean (mm plant^-1^) | 12.84 ** | 3.41 ns | 0.03 ns |
|  | | Root diameter maximum (mm plant^-1^) | 3.36 ns | 7.02 * | 4.47 ns |
|  | | Root diameter median (mm plant^-1^) | 0.21 ns | 0.24 ns | 0.00 ns |
| Root respiration | | Root CO_2_ flux total (nmol plant^-1^ s^-1^) | 8.39 * | 0.18 ns | 0.74 ns |
|  | | Specific root CO_2_ flux (nmol g^-1^ s^-1^) | 7.74 * | 0.02 ns | 0.29 ns |
|  | | Specific root CO_2_ flux (nmol m^-1^ s^-1^) | 2.34 ns | 0.10 ns | 0.18 ns |
| Biomass distribution | | Root mass fraction (g g^1^) | 21.69 *** | 4.08 ns | 0.13 ns |
|  | | Total plant mass (g plant^-1^) | 5.30 * | 6.13 * | 0.41 ns |
| Shoot size | | Shoot dry mass total (g plant^-1^) | 0.82 ns | 8.45 * | 0.32 ns |
|  | | Plant height (cm plant^-1^) | 0.05 ns | 3.90 ns | 0.99 ns |
|  | | Tiller count | 4.90 * | 6.60 * | 4.15 ns |
|  | | Leaf maximum width (cm) | 12.12 ** | 6.19 * | 0.00 ns |
| Shoot properties | | Shoot carbon content (%) | 0.34 ns | 60.52 *** | 1.06 ns |
|  | | Shoot N concentration (%) | 4.42 ns | 2.16 ns | 0.62 ns |
|  | | Shoot 15N concentration (%) | 0.01 ns | 3.63 ns | 0.07 ns |
|  | | Shoot total 15N content (mg plant^-1^) | 0.44 ns | 8.54 ** | 1.47 ns |
|  | | Shoot 15N uptake rate (mg plant^-1^ h^-1^) | 0.44 ns | 8.54 ** | 1.47 ns |
|  | | CO_2_ assimilation rate (µmol m^-2^ s^-1^) | 0.14 ns | 0.22 ns | 0.09 ns |
|  | | Transpiration rate (mol m^-2^ s^-1^) | 0.40 ns | 1.09 ns | 2.30 ns |
|  | | Stomatal conductance (mol m^-2^ s^-1^) | 0.44 ns | 0.79 ns | 2.75 ns |
|  | | Intracellular CO_2_ (Pci) | 0.02 ns | 0.40 ns | 0.09 ns |

*** P < 0.001; ** P < 0.01; * P < 0.05; ns not significant

Table S6. Genotype x W condition (low N conditions)

|  | Traits | Source of variation | | |
| --- | --- | --- | --- | --- |
|  |  | Geno | W Treatment | Geno:W Treatment |
| Total root size | Root dry mass total (g plant^-1^) | 14.15 ** | 0.97 ns | 0.50 ns |
|  | Root length total (mm plant^-1^) | 7.52 * | 2.60 ns | 1.17 ns |
|  | Root length axial (mm plant^-1^) | 17.73 *** | 1.55 ns | 0.01 ns |
|  | Root length lateral (mm plant^-1^) | 4.42 ns | 2.54 ns | 1.10 ns |
|  | Root length secondary lateral (mm plant^-1^) | 7.98 * | 2.40 ns | 1.34 ns |
|  | Root surface area total (mm^2^ plant^-1^) | 10.22 ** | 2.82 ns | 1.00 ns |
|  | Root surface area axial (mm^2^ plant^-1^) | 18.72 *** | 1.11 ns | 0.03 ns |
|  | Root surface area lateral (mm^2^ plant^-1^) | 4.33 ns | 2.66 ns | 1.17 ns |
|  | Root surface area secondary lateral (mm^2^ plant^-1^) | 8.23 * | 2.66 ns | 1.37 ns |
|  | Root volume total (mm^3^ plant^-1^) | 16.18 *** | 2.24 ns | 0.63 ns |
|  | Root volume axial (mm^3^ plant^-1^) | 17.15 *** | 0.64 ns | 0.06 ns |
|  | Root volume lateral (mm^3^ plant^-1^) | 4.36 ns | 2.80 ns | 1.15 ns |
|  | Root volume secondary lateral (mm^3^ plant^-1^) | 8.38 * | 2.83 ns | 1.37 ns |
|  | Root branch count total | 7.16 * | 1.23 ns | 0.51 ns |
|  | Root tip count total | 8.43 * | 1.85 ns | 1.16 ns |
| Root distribution | Specific root length (m g^-1^) | 8.02 * | 1.44 ns | 0.12 ns |
|  | Root lateral:axial root ratio (ratio) | 10.09 ** | 0.06 ns | 0.73 ns |
|  | Root branching frequency (branch mm^-1^) | 1.74 ns | 0.08 ns | 0.00 ns |
|  | Deep root mass total (g plant^-1^) | 17.73 *** | 1.23 ns | 1.63 ns |
|  | Deep root length total (mm plant^-1^) | 9.54 ** | 3.53 ns | 2.28 ns |
|  | Deep root mass fraction (g g^1^) | 3.99 ns | 0.27 ns | 0.11 ns |
|  | Deep root length fraction (mm mm^1^) | 5.84 * | 1.54 ns | 0.43 ns |
| Root diameter | Root diameter mean (mm plant^-1^) | 9.00 ** | 0.09 ns | 1.06 ns |
|  | Root diameter maximum (mm plant^-1^) | 14.01 ** | 0.00 ns | 0.04 ns |
|  | Root diameter median (mm plant^-1^) | 0.30 ns | 1.32 ns | 0.54 ns |
| Root respiration | Root CO_2_ flux total (nmol plant^-1^ s^-1^) | 4.50 ns | 0.95 ns | 0.64 ns |
|  | Specific root CO_2_ flux (nmol g^-1^ s^-1^) | 8.47 * | 1.37 ns | 2.54 ns |
|  | Specific root CO_2_ flux (nmol m^-1^ s^-1^) | 2.43 ns | 0.51 ns | 2.92 ns |
| Biomass distribution | Root mass fraction (g g^1^) | 10.22 ** | 0.33 ns | 0.20 ns |
|  | Total plant mass (g plant^-1^) | 8.25 * | 1.56 ns | 0.30 ns |
| Shoot size | Shoot dry mass total (g plant^-1^) | 3.20 ns | 1.67 ns | 0.12 ns |
|  | Plant height (cm plant^-1^) | 18.66 *** | 4.35 ns | 0.30 ns |
|  | Tiller count | 0.63 ns | 0.16 ns | 0.63 ns |
|  | Leaf maximum width (cm) | 8.11 * | 1.20 ns | 0.05 ns |
| Shoot properties | Shoot carbon content (%) | 6.87 * | 10.61 ** | 0.77 ns |
|  | Shoot N concentration (%) | 0.04 ns | 0.04 ns | 0.67 ns |
|  | Shoot 15N concentration (%) | 1.34 ns | 1.07 ns | 0.43 ns |
|  | Shoot total 15N content (mg plant^-1^) | 5.95 * | 0.14 ns | 0.20 ns |
|  | Shoot 15N uptake rate (mg plant^-1^ h^-1^) | 5.95 * | 0.14 ns | 0.20 ns |
|  | CO_2_ assimilation rate (µmol m^-2^ s^-1^) | 1.97 ns | 0.01 ns | 0.17 ns |
|  | Transpiration rate (mol m^-2^ s^-1^) | 1.14 ns | 1.19 ns | 0.06 ns |
|  | Stomatal conductance (mol m^-2^ s^-1^) | 1.13 ns | 1.19 ns | 0.14 ns |
|  | Intracellular CO_2_ (Pci) | 1.06 ns | 0.00 ns | 0.05 ns |

*** P < 0.001; ** P < 0.01; * P < 0.05; ns not significant

Table S7. Genotype x Depth (HNWW conditions)

|  | Traits | Source of variation | | |
| --- | --- | --- | --- | --- |
|  |  | Geno | Depth | Geno:Depth |
| Total root size | Root dry mass (g plant^-1^) | 14.99 *** | 17.59 *** | 0.56 ns |
|  | Root length total (mm plant^-1^) | 13.47 *** | 1.74 ns | 0.45 ns |
|  | Root length axial (mm plant^-1^) | 7.18 *** | 2.87 ns | 0.28 ns |
|  | Root length lateral (mm plant^-1^) | 11.05 *** | 4.69 * | 0.77 ns |
|  | Root length secondary lateral (mm plant^-1^) | 16.90 *** | 1.01 ns | 0.49 ns |
|  | Root surface area total (mm^2^ plant^-1^) | 11.26 *** | 0.18 ns | 0.12 ns |
|  | Root surface area axial (mm^2^ plant^-1^) | 7.99 *** | 6.99 * | 0.36 ns |
|  | Root surface area lateral (mm^2^ plant^-1^) | 10.72 *** | 5.60 * | 0.73 ns |
|  | Root surface area secondary lateral (mm^2^ plant^-1^) | 15.66 *** | 0.93 ns | 0.44 ns |
|  | Root volume total (mm^3^ plant^-1^) | 10.09 *** | 2.15 ns | 0.02 ns |
|  | Root volume axial (mm^3^ plant^-1^) | 8.81 *** | 11.84 ** | 0.43 ns |
|  | Root volume lateral (mm^3^ plant^-1^) | 10.40 *** | 6.36 * | 0.67 ns |
|  | Root volume secondary lateral (mm^3^ plant^-1^) | 14.93 *** | 0.87 ns | 0.41 ns |
|  | Root branch count total | 19.50 *** | 5.93 * | 0.70 ns |
|  | Root tip count total | 15.87 *** | 2.02 ns | 0.85 ns |
| Root distribution | Specific root length (m g^-1^) | 11.43 *** | 24.81 *** | 0.88 ns |
|  | Root lateral:axial root ratio (ratio) | 5.07 ** | 14.04 *** | 2.20 ns |
|  | Root branching frequency (branch mm^-1^) | 70.38 *** | 19.01 *** | 0.63 ns |
| Root diameter | Root diameter mean (mm plant^-1^) | 16.82 *** | 34.32 *** | 2.61 ns |
|  | Root diameter maximum (mm plant^-1^) | 2.78 * | 2.23 ns | 0.91 ns |
|  | Root diameter median (mm plant^-1^) | 13.59 *** | 1.74 ns | 0.90 ns |
| Root respiration | Specific root CO_2_ flux (nmol g^-1^ s^-1^) | 1.94 ns | 44.89 *** | 2.92 * |
|  | Specific root CO_2_ flux (nmol m^-1^ s^-1^) | 0.76 ns | 2.95 ns | 1.27 ns |

*** P < 0.001; ** P < 0.01; * P < 0.05; ns not significant

Table S8. Genotype x Depth (LNWW conditions)

|  | Traits | Source of variation | | |
| --- | --- | --- | --- | --- |
|  |  | Geno | Depth | Geno:Depth |
| Total root size | Root dry mass (g plant^-1^) | 12.88 *** | 16.13 *** | 0.46 ns |
|  | Root length total (mm plant^-1^) | 8.75 *** | 7.62 ** | 2.55 ns |
|  | Root length axial (mm plant^-1^) | 20.23 *** | 21.97 *** | 0.50 ns |
|  | Root length lateral (mm plant^-1^) | 8.28 *** | 6.26 * | 2.65 * |
|  | Root length secondary lateral (mm plant^-1^) | 8.18 *** | 6.85 * | 2.56 ns |
|  | Root surface area total (mm^2^ plant^-1^) | 9.60 *** | 10.26 ** | 2.12 ns |
|  | Root surface area axial (mm^2^ plant^-1^) | 22.86 *** | 26.55 *** | 0.73 ns |
|  | Root surface area lateral (mm^2^ plant^-1^) | 8.53 *** | 6.28 * | 2.66 * |
|  | Root surface area secondary lateral (mm^2^ plant^-1^) | 7.70 *** | 7.37 ** | 2.50 ns |
|  | Root volume total (mm^3^ plant^-1^) | 12.74 *** | 16.73 *** | 1.31 ns |
|  | Root volume axial (mm^3^ plant^-1^) | 23.61 *** | 29.14 *** | 1.08 ns |
|  | Root volume lateral (mm^3^ plant^-1^) | 8.53 *** | 6.39 * | 2.54 ns |
|  | Root volume secondary lateral (mm^3^ plant^-1^) | 7.37 *** | 7.76 ** | 2.46 ns |
|  | Root branch count total | 11.30 *** | 5.15 * | 2.49 ns |
|  | Root tip count total | 10.24 *** | 8.15 ** | 2.28 ns |
| Root distribution | Specific root length (m g^-1^) | 4.87 ** | 9.74 ** | 0.96 ns |
|  | Root lateral:axial root ratio (ratio) | 1.18 ns | 4.70 * | 1.28 ns |
|  | Root branching frequency (branch mm^-1^) | 15.79 *** | 2.29 ns | 0.10 ns |
| Root diameter | Root diameter mean (mm plant^-1^) | 1.84 ns | 7.38 * | 0.99 ns |
|  | Root diameter maximum (mm plant^-1^) | 5.82 ** | 15.60 *** | 0.11 ns |
|  | Root diameter median (mm plant^-1^) | 2.08 ns | 0.00 ns | 0.63 ns |
| Root respiration | Specific root CO_2_ flux (nmol g^-1^ s^-1^) | 1.41 ns | 8.10 ** | 0.79 ns |
|  | Specific root CO_2_ flux (nmol m^-1^ s^-1^) | 1.40 ns | 7.68 * | 0.38 ns |

*** P < 0.001; ** P < 0.01; * P < 0.05; ns not significant

Table S9. Genotype x Depth (HNDS conditions)

|  | Traits | Source of variation | | |
| --- | --- | --- | --- | --- |
|  |  | Geno | Depth | Geno:Depth |
| Total root size | Root dry mass (g plant^-1^) | 10.65 *** | 66.52 *** | 0.70 ns |
|  | Root length total (mm plant^-1^) | 28.56 *** | 0.01 ns | 0.99 ns |
|  | Root length axial (mm plant^-1^) | 22.42 *** | 38.21 *** | 2.11 ns |
|  | Root length lateral (mm plant^-1^) | 16.88 *** | 1.75 ns | 0.87 ns |
|  | Root length secondary lateral (mm plant^-1^) | 35.93 *** | 0.24 ns | 0.88 ns |
|  | Root surface area total (mm^2^ plant^-1^) | 25.61 *** | 0.91 ns | 1.25 ns |
|  | Root surface area axial (mm^2^ plant^-1^) | 20.41 *** | 63.06 *** | 1.56 ns |
|  | Root surface area lateral (mm^2^ plant^-1^) | 17.33 *** | 2.95 ns | 1.01 ns |
|  | Root surface area secondary lateral (mm^2^ plant^-1^) | 34.02 *** | 0.22 ns | 0.90 ns |
|  | Root volume total (mm^3^ plant^-1^) | 26.77 *** | 20.61 *** | 1.09 ns |
|  | Root volume axial (mm^3^ plant^-1^) | 16.65 *** | 75.58 *** | 0.79 ns |
|  | Root volume lateral (mm^3^ plant^-1^) | 17.93 *** | 4.43 * | 1.19 ns |
|  | Root volume secondary lateral (mm^3^ plant^-1^) | 32.08 *** | 0.21 ns | 0.90 ns |
|  | Root branch count total | 36.95 *** | 0.16 ns | 0.72 ns |
|  | Root tip count total | 30.44 *** | 1.58 ns | 1.97 ns |
| Root distribution | Specific root length (m g^-1^) | 20.79 *** | 27.77 *** | 1.68 ns |
|  | Root lateral:axial root ratio (ratio) | 30.01 *** | 17.67 *** | 1.18 ns |
|  | Root branching frequency (branch mm^-1^) | 110.23 *** | 0.01 ns | 0.63 ns |
| Root diameter | Root diameter mean (mm plant^-1^) | 25.17 *** | 21.99 *** | 1.63 ns |
|  | Root diameter maximum (mm plant^-1^) | 6.22 *** | 10.30 ** | 4.27 ** |
|  | Root diameter median (mm plant^-1^) | 10.13 *** | 0.91 ns | 0.24 ns |
| Root respiration | Specific root CO_2_ flux (nmol g^-1^ s^-1^) | 0.56 ns | 7.09 * | 0.55 ns |
|  | Specific root CO_2_ flux (nmol m^-1^ s^-1^) | 0.22 ns | 2.67 ns | 0.64 ns |

*** P < 0.001; ** P < 0.01; * P < 0.05; ns not significant

Table S10. Genotype x Depth (LNDS conditions)

|  | Traits | Source of variation | | |
| --- | --- | --- | --- | --- |
|  |  | Geno | Depth | Geno:Depth |
| Total root size | Root dry mass (g plant^-1^) | 42.18 *** | 26.92 *** | 1.08 ns |
|  | Root length total (mm plant^-1^) | 23.14 *** | 4.44 * | 1.59 ns |
|  | Root length axial (mm plant^-1^) | 33.12 *** | 45.52 *** | 2.96 * |
|  | Root length lateral (mm plant^-1^) | 15.41 *** | 2.26 ns | 1.32 ns |
|  | Root length secondary lateral (mm plant^-1^) | 25.76 *** | 3.54 ns | 1.71 ns |
|  | Root surface area total (mm^2^ plant^-1^) | 26.39 *** | 9.61 ** | 1.83 ns |
|  | Root surface area axial (mm^2^ plant^-1^) | 33.98 *** | 47.91 *** | 2.78 * |
|  | Root surface area lateral (mm^2^ plant^-1^) | 15.44 *** | 2.01 ns | 1.28 ns |
|  | Root surface area secondary lateral (mm^2^ plant^-1^) | 22.51 *** | 3.74 ns | 1.63 ns |
|  | Root volume total (mm^3^ plant^-1^) | 35.23 *** | 26.14 *** | 2.27 ns |
|  | Root volume axial (mm^3^ plant^-1^) | 28.72 *** | 41.28 *** | 2.04 ns |
|  | Root volume lateral (mm^3^ plant^-1^) | 14.85 *** | 2.03 ns | 1.16 ns |
|  | Root volume secondary lateral (mm^3^ plant^-1^) | 20.34 *** | 3.91 ns | 1.57 ns |
|  | Root branch count total | 38.75 *** | 4.93 * | 2.25 ns |
|  | Root tip count total | 23.74 *** | 4.90 * | 1.47 ns |
| Root distribution | Specific root length (m g^-1^) | 15.51 *** | 44.68 *** | 2.50 ns |
|  | Root lateral:axial root ratio (ratio) | 0.89 ns | 2.67 ns | 0.91 ns |
|  | Root branching frequency (branch mm^-1^) | 33.31 *** | 0.77 ns | 2.06 ns |
| Root diameter | Root diameter mean (mm plant^-1^) | 4.39 ** | 31.80 *** | 2.69 * |
|  | Root diameter maximum (mm plant^-1^) | 6.39 *** | 14.91 *** | 1.90 ns |
|  | Root diameter median (mm plant^-1^) | 8.52 *** | 2.31 ns | 1.09 ns |
| Root respiration | Specific root CO_2_ flux (nmol g^-1^ s^-1^) | 5.56 ** | 20.81 *** | 0.82 ns |
|  | Specific root CO_2_ flux (nmol m^-1^ s^-1^) | 9.61 *** | 0.01 ns | 0.38 ns |

*** P < 0.001; ** P < 0.01; * P < 0.05; ns not significant

Table S11. NTreatment x WTreatment (both ecotypes)

|  | Traits | Source of variation | | |
| --- | --- | --- | --- | --- |
|  |  | N Treat | W Treat | N Treat:W Treat |
| Total root size | Root dry mass total (g plant^-1^) | 75.02 *** | 1.25 ns | 0.69 ns |
|  | Root length total (mm plant^-1^) | 85.10 *** | 3.97 ns | 0.64 ns |
|  | Root length axial (mm plant^-1^) | 119.47 *** | 13.13 *** | 10.01 ** |
|  | Root length lateral (mm plant^-1^) | 62.18 *** | 6.73 * | 3.00 ns |
|  | Root length secondary lateral (mm plant^-1^) | 48.07 *** | 0.04 ns | 2.77 ns |
|  | Root surface area total (mm^2^ plant^-1^) | 127.79 *** | 11.91 ** | 6.40 * |
|  | Root surface area axial (mm^2^ plant^-1^) | 119.28 *** | 13.19 *** | 10.90 ** |
|  | Root surface area lateral (mm^2^ plant^-1^) | 60.91 *** | 6.82 * | 3.30 ns |
|  | Root surface area secondary lateral (mm^2^ plant^-1^) | 48.98 *** | 0.20 ns | 1.28 ns |
|  | Root volume total (mm^3^ plant^-1^) | 154.84 *** | 17.23 *** | 13.08 *** |
|  | Root volume axial (mm^3^ plant^-1^) | 110.74 *** | 12.59 ** | 11.13 ** |
|  | Root volume lateral (mm^3^ plant^-1^) | 59.16 *** | 6.88 * | 3.51 ns |
|  | Root volume secondary lateral (mm^3^ plant^-1^) | 49.60 *** | 0.84 ns | 0.54 ns |
|  | Root branch count total | 56.71 *** | 0.17 ns | 0.05 ns |
|  | Root tip count total | 112.60 *** | 6.64 * | 2.61 ns |
| Root distribution | Specific root length (m g^-1^) | 38.11 *** | 0.43 ns | 1.00 ns |
|  | Root lateral:axial root ratio (ratio) | 33.92 *** | 0.30 ns | 0.84 ns |
|  | Root branching frequency (branch mm^-1^) | 0.02 ns | 0.28 ns | 0.01 ns |
|  | Deep root mass total (g plant^-1^) | 90.11 *** | 0.38 ns | 1.10 ns |
|  | Deep root length total (mm plant^-1^) | 22.27 *** | 3.95 ns | 0.29 ns |
|  | Deep root mass fraction (g g^1^) | 3.00 ns | 2.31 ns | 5.29 * |
|  | Deep root length fraction (mm mm^1^) | 0.26 ns | 1.87 ns | 0.03 ns |
| Root diameter | Root diameter mean (mm plant^-1^) | 102.59 *** | 1.28 ns | 1.91 ns |
|  | Root diameter maximum (mm plant^-1^) | 58.09 *** | 1.09 ns | 1.13 ns |
|  | Root diameter median (mm plant^-1^) | 108.52 *** | 0.01 ns | 0.74 ns |
| Root respiration | Root CO_2_ flux total (nmol plant^-1^ s^-1^) | 5.57 * | 0.22 ns | 0.04 ns |
|  | Specific root CO_2_ flux (nmol g^-1^ s^-1^) | 9.89 ** | 1.40 ns | 1.56 ns |
|  | Specific root CO_2_ flux (nmol m^-1^ s^-1^) | 1.25 ns | 0.39 ns | 0.81 ns |
| Biomass distribution | Root mass fraction (g g^1^) | 0.39 ns | 1.52 ns | 0.25 ns |
|  | Total plant mass (g plant^-1^) | 146.09 *** | 5.84 * | 4.03 ns |
| Shoot size | Shoot dry mass total (g plant^-1^) | 175.36 *** | 10.40 ** | 7.69 ** |
|  | Plant height (cm plant^-1^) | 94.07 *** | 6.25 * | 1.13 ns |
|  | Tiller count | 49.23 *** | 4.64 * | 3.52 ns |
|  | Leaf maximum width (cm) | 62.54 *** | 0.42 ns | 3.78 ns |
| Shoot properties | Shoot carbon content (%) | 59.84 *** | 26.63 *** | 0.05 ns |
|  | Shoot N concentration (%) | 18.04 *** | 1.03 ns | 1.56 ns |
|  | Shoot 15N concentration (%) | 9.49 ** | 1.02 ns | 1.14 ns |
|  | Shoot total 15N content (mg plant^-1^) | 82.85 *** | 7.36 * | 8.48 ** |
|  | Shoot 15N uptake rate (mg plant^-1^ h^-1^) | 82.85 *** | 7.36 * | 8.48 ** |
|  | CO_2_ assimilation rate (µmol m^-2^ s^-1^) | 0.10 ns | 0.03 ns | 0.13 ns |
|  | Transpiration rate (mol m^-2^ s^-1^) | 0.78 ns | 2.24 ns | 0.03 ns |
|  | Stomatal conductance (mol m^-2^ s^-1^) | 0.70 ns | 1.81 ns | 0.00 ns |
|  | Intracellular CO_2_ (Pci) | 0.17 ns | 0.19 ns | 0.13 ns |

*** P < 0.001; ** P < 0.01; * P < 0.05; ns not significant

Table S12. NTreatment x WTreatment (AP13)

|  | Traits | Source of variation | | |
| --- | --- | --- | --- | --- |
|  |  | N Treat | W Treat | N Treat:W Treat |
| Total root size | Root dry mass total (g plant^-1^) | 90.55 *** | 4.85 * | 4.33 ns |
|  | Root length total (mm plant^-1^) | 50.73 *** | 2.24 ns | 1.53 ns |
|  | Root length axial (mm plant^-1^) | 54.55 *** | 8.50 * | 6.76 * |
|  | Root length lateral (mm plant^-1^) | 60.64 *** | 4.92 * | 3.94 ns |
|  | Root length secondary lateral (mm plant^-1^) | 26.72 *** | 0.00 ns | 0.04 ns |
|  | Root surface area total (mm^2^ plant^-1^) | 63.57 *** | 6.00 * | 4.85 * |
|  | Root surface area axial (mm^2^ plant^-1^) | 62.14 *** | 10.54 ** | 9.09 * |
|  | Root surface area lateral (mm^2^ plant^-1^) | 60.49 *** | 4.94 * | 4.05 ns |
|  | Root surface area secondary lateral (mm^2^ plant^-1^) | 29.11 *** | 0.18 ns | 0.02 ns |
|  | Root volume total (mm^3^ plant^-1^) | 71.60 *** | 10.29 ** | 9.14 * |
|  | Root volume axial (mm^3^ plant^-1^) | 68.40 *** | 13.03 ** | 12.01 ** |
|  | Root volume lateral (mm^3^ plant^-1^) | 58.86 *** | 4.87 * | 4.00 ns |
|  | Root volume secondary lateral (mm^3^ plant^-1^) | 30.93 *** | 0.50 ns | 0.15 ns |
|  | Root branch count total | 29.40 *** | 0.71 ns | 0.50 ns |
|  | Root tip count total | 68.76 *** | 2.30 ns | 1.93 ns |
| Root distribution | Specific root length (m g^-1^) | 22.23 *** | 0.43 ns | 0.91 ns |
|  | Root lateral:axial root ratio (ratio) | 28.37 *** | 0.95 ns | 0.08 ns |
|  | Root branching frequency (branch mm^-1^) | 0.54 ns | 0.11 ns | 0.37 ns |
|  | Deep root mass total (g plant^-1^) | 39.11 *** | 0.01 ns | 0.02 ns |
|  | Deep root length total (mm plant^-1^) | 20.02 *** | 0.20 ns | 0.06 ns |
|  | Deep root mass fraction (g g^1^) | 6.26 * | 1.95 ns | 2.51 ns |
|  | Deep root length fraction (mm mm^1^) | 0.74 ns | 0.14 ns | 0.98 ns |
| Root diameter | Root diameter mean (mm plant^-1^) | 109.74 *** | 2.84 ns | 0.92 ns |
|  | Root diameter maximum (mm plant^-1^) | 60.80 *** | 2.11 ns | 2.83 ns |
|  | Root diameter median (mm plant^-1^) | 93.70 *** | 0.15 ns | 0.41 ns |
| Root respiration | Root CO_2_ flux total (nmol plant^-1^ s^-1^) | 8.01 * | 0.49 ns | 0.41 ns |
|  | Specific root CO_2_ flux (nmol g^-1^ s^-1^) | 8.85 ** | 1.74 ns | 2.32 ns |
|  | Specific root CO_2_ flux (nmol m^-1^ s^-1^) | 1.87 ns | 0.91 ns | 1.95 ns |
| Biomass distribution | Root mass fraction (g g^1^) | 0.39 ns | 2.04 ns | 0.00 ns |
|  | Total plant mass (g plant^-1^) | 94.45 *** | 7.20 * | 6.10 * |
| Shoot size | Shoot dry mass total (g plant^-1^) | 87.73 *** | 7.74 * | 6.44 * |
|  | Plant height (cm plant^-1^) | 43.91 *** | 3.74 ns | 1.68 ns |
|  | Tiller count | 34.24 *** | 7.74 * | 4.87 * |
|  | Leaf maximum width (cm) | 35.76 *** | 0.16 ns | 2.54 ns |
| Shoot properties | Shoot carbon content (%) | 23.36 *** | 25.20 *** | 0.00 ns |
|  | Shoot N concentration (%) | 38.71 *** | 1.54 ns | 6.34 * |
|  | Shoot 15N concentration (%) | 1.65 ns | 1.26 ns | 1.33 ns |
|  | Shoot total 15N content (mg plant^-1^) | 75.30 *** | 9.83 ** | 12.29 ** |
|  | Shoot 15N uptake rate (mg plant^-1^ h^-1^) | 75.30 *** | 9.83 ** | 12.29 ** |
|  | CO_2_ assimilation rate (µmol m^-2^ s^-1^) | 0.77 ns | 0.00 ns | 0.51 ns |
|  | Transpiration rate (mol m^-2^ s^-1^) | 0.03 ns | 0.01 ns | 0.34 ns |
|  | Stomatal conductance (mol m^-2^ s^-1^) | 0.04 ns | 0.01 ns | 0.44 ns |
|  | Intracellular CO_2_ (Pci) | 0.16 ns | 0.11 ns | 0.33 ns |

*** P < 0.001; ** P < 0.01; * P < 0.05; ns not significant

Table S13. NTreatment x WTreatment (VS16)

|  | Traits | Source of variation | | |
| --- | --- | --- | --- | --- |
|  |  | N Treat | W Treat | N Treat:W Treat |
| Total root size | Root dry mass total (g plant^-1^) | 67.50 *** | 0.33 ns | 0.04 ns |
|  | Root length total (mm plant^-1^) | 34.40 *** | 1.73 ns | 0.05 ns |
|  | Root length axial (mm plant^-1^) | 86.85 *** | 6.76 * | 4.91 * |
|  | Root length lateral (mm plant^-1^) | 15.61 ** | 2.53 ns | 0.40 ns |
|  | Root length secondary lateral (mm plant^-1^) | 25.17 *** | 0.19 ns | 7.55 * |
|  | Root surface area total (mm^2^ plant^-1^) | 57.75 *** | 5.30 * | 1.54 ns |
|  | Root surface area axial (mm^2^ plant^-1^) | 100.54 *** | 7.56 * | 5.96 * |
|  | Root surface area lateral (mm^2^ plant^-1^) | 15.44 ** | 2.72 ns | 0.55 ns |
|  | Root surface area secondary lateral (mm^2^ plant^-1^) | 23.38 *** | 0.04 ns | 4.81 * |
|  | Root volume total (mm^3^ plant^-1^) | 101.18 *** | 8.79 ** | 5.52 * |
|  | Root volume axial (mm^3^ plant^-1^) | 103.67 *** | 7.63 * | 6.44 * |
|  | Root volume lateral (mm^3^ plant^-1^) | 15.34 ** | 2.93 ns | 0.70 ns |
|  | Root volume secondary lateral (mm^3^ plant^-1^) | 21.96 *** | 0.40 ns | 3.14 ns |
|  | Root branch count total | 33.49 *** | 0.31 ns | 2.39 ns |
|  | Root tip count total | 54.98 *** | 6.28 * | 0.90 ns |
| Root distribution | Specific root length (m g^-1^) | 42.07 *** | 0.22 ns | 0.60 ns |
|  | Root lateral:axial root ratio (ratio) | 24.67 *** | 0.08 ns | 2.83 ns |
|  | Root branching frequency (branch mm^-1^) | 0.97 ns | 1.24 ns | 0.62 ns |
|  | Deep root mass total (g plant^-1^) | 100.41 *** | 1.51 ns | 4.32 ns |
|  | Deep root length total (mm plant^-1^) | 5.08 * | 6.21 * | 0.29 ns |
|  | Deep root mass fraction (g g^1^) | 0.05 ns | 0.53 ns | 2.96 ns |
|  | Deep root length fraction (mm mm^1^) | 2.05 ns | 5.01 * | 0.32 ns |
| Root diameter | Root diameter mean (mm plant^-1^) | 72.34 *** | 0.39 ns | 2.13 ns |
|  | Root diameter maximum (mm plant^-1^) | 28.99 *** | 0.10 ns | 0.01 ns |
|  | Root diameter median (mm plant^-1^) | 37.98 *** | 0.01 ns | 0.37 ns |
| Root respiration | Root CO_2_ flux total (nmol plant^-1^ s^-1^) | 1.22 ns | 0.06 ns | 1.92 ns |
|  | Specific root CO_2_ flux (nmol g^-1^ s^-1^) | 20.54 *** | 0.80 ns | 2.42 ns |
|  | Specific root CO_2_ flux (nmol m^-1^ s^-1^) | 0.05 ns | 2.40 ns | 2.87 ns |
| Biomass distribution | Root mass fraction (g g^1^) | 0.38 ns | 1.06 ns | 0.77 ns |
|  | Total plant mass (g plant^-1^) | 87.53 *** | 1.75 ns | 0.85 ns |
| Shoot size | Shoot dry mass total (g plant^-1^) | 75.48 *** | 2.86 ns | 1.77 ns |
|  | Plant height (cm plant^-1^) | 72.51 *** | 3.30 ns | 0.00 ns |
|  | Tiller count | 39.02 *** | 0.10 ns | 0.39 ns |
|  | Leaf maximum width (cm) | 62.00 *** | 0.58 ns | 3.16 ns |
| Shoot properties | Shoot carbon content (%) | 40.72 *** | 7.70 * | 0.13 ns |
|  | Shoot N concentration (%) | 3.88 ns | 0.40 ns | 0.01 ns |
|  | Shoot 15N concentration (%) | 9.89 ** | 0.07 ns | 0.09 ns |
|  | Shoot total 15N content (mg plant^-1^) | 24.24 *** | 1.11 ns | 1.05 ns |
|  | Shoot 15N uptake rate (mg plant^-1^ h^-1^) | 24.24 *** | 1.11 ns | 1.05 ns |
|  | CO_2_ assimilation rate (µmol m^-2^ s^-1^) | 1.52 ns | 0.06 ns | 0.01 ns |
|  | Transpiration rate (mol m^-2^ s^-1^) | 2.46 ns | 4.84 * | 0.88 ns |
|  | Stomatal conductance (mol m^-2^ s^-1^) | 2.43 ns | 5.05 * | 0.78 ns |
|  | Intracellular CO_2_ (Pci) | 0.93 ns | 0.10 ns | 0.00 ns |

*** P < 0.001; ** P < 0.01; * P < 0.05; ns not significant

*Other data probably wont use*

### 1.1 [ Seedling growth study ] (Need to decide on root class names, here seminal root system)

Growing from seed in hydroponics, the upland ecotype (Summer) had significantly greater root length and shoot size in the seedling study,

Significant differences for root traits were observed from 9 days, with the difference becoming greater with age (d9 – d30)

Differences observed in roots appeared to be across crown and lateral roots, and not secondary lateral roots, and a difference in seedling vigor is possible. Lateral roots were the greatest difference with significant differences at all ages. Crown root differences were from d23 and d30.

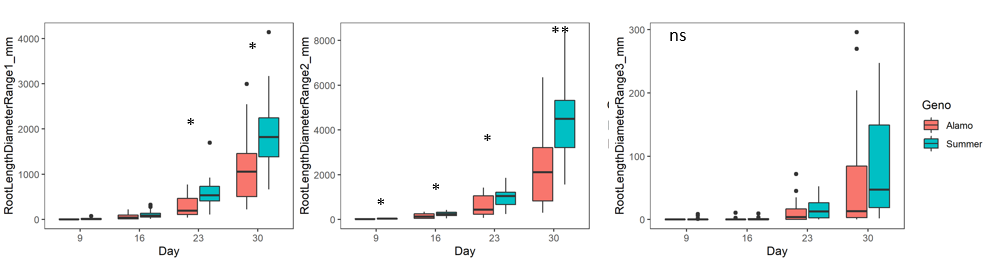

Significant differences for shoot traits were observed from 23 days and 30 days

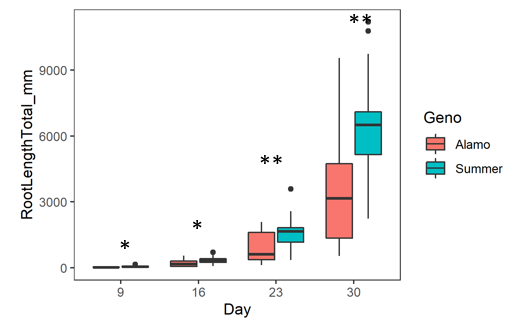

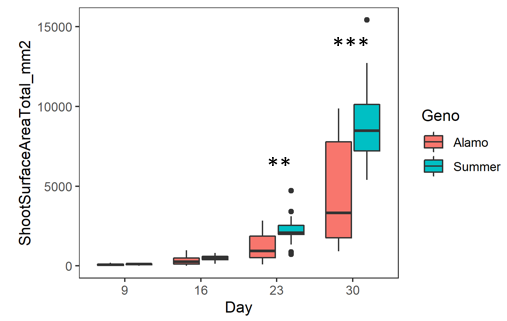

### 1.2 [ Seedling nutrient uptake study ]
